# *FKS1/2*-variant independent mechanisms underlying the emergence of resistance in echinocandin-refractory *Candida auris* infections

**DOI:** 10.64898/2026.01.17.700071

**Authors:** Hugh Gifford, Arnab Pradhan, Ben Caswall, Sarah Murphy, Ian Leaves, Matt Edmondson, Tanmoy Chakraborty, Duncan Wilson, Surabhi K. Taori, Neil A. R. Gow, Rhys A. Farrer, Alistair J. P. Brown, Tihana Bicanic

## Abstract

The emerging fungus *Candida auris* is a drug resistant global public health threat and WHO critical priority pathogen. Recommended first-line invasive candidiasis treatment is echinocandin monotherapy, but *C. auris* can develop on-treatment resistance *via FKS1/2* gene mutations and additional, previously unexplained mechanisms. To better understand echinocandin failure in *C. auris*, we sequenced the genomes of echinocandin refractory *FKS1/2* wild-type *C. auris* serial isolates from two critically unwell patients in London, UK. Population analysis profiling revealed echinocandin heteroresistance, and *in vitro* culture of clinical isolates at supra-MIC concentrations of anidulafungin (8 μg/ml) exhibited morphotypic heterogeneity. Small colony variants (SCVs) and large colony variants (LCVs) showed elevated MICs with polyploidy (to 4n and above) alongside adaptive changes in cell wall β-1,3-glucan content. LCVs contained significantly more mutations in calcineurin-related stress tolerance pathway gene *CRZ1* compared to clinical parents and SCVs, associated with further increases in MIC. These findings indicate progressive step-wise accrual of adaptation to echinocandins, including genomic instability, alterations in stress tolerance pathways, and cell wall remodeling, paving the way for resistance emergence.

**Graphical abstract:** 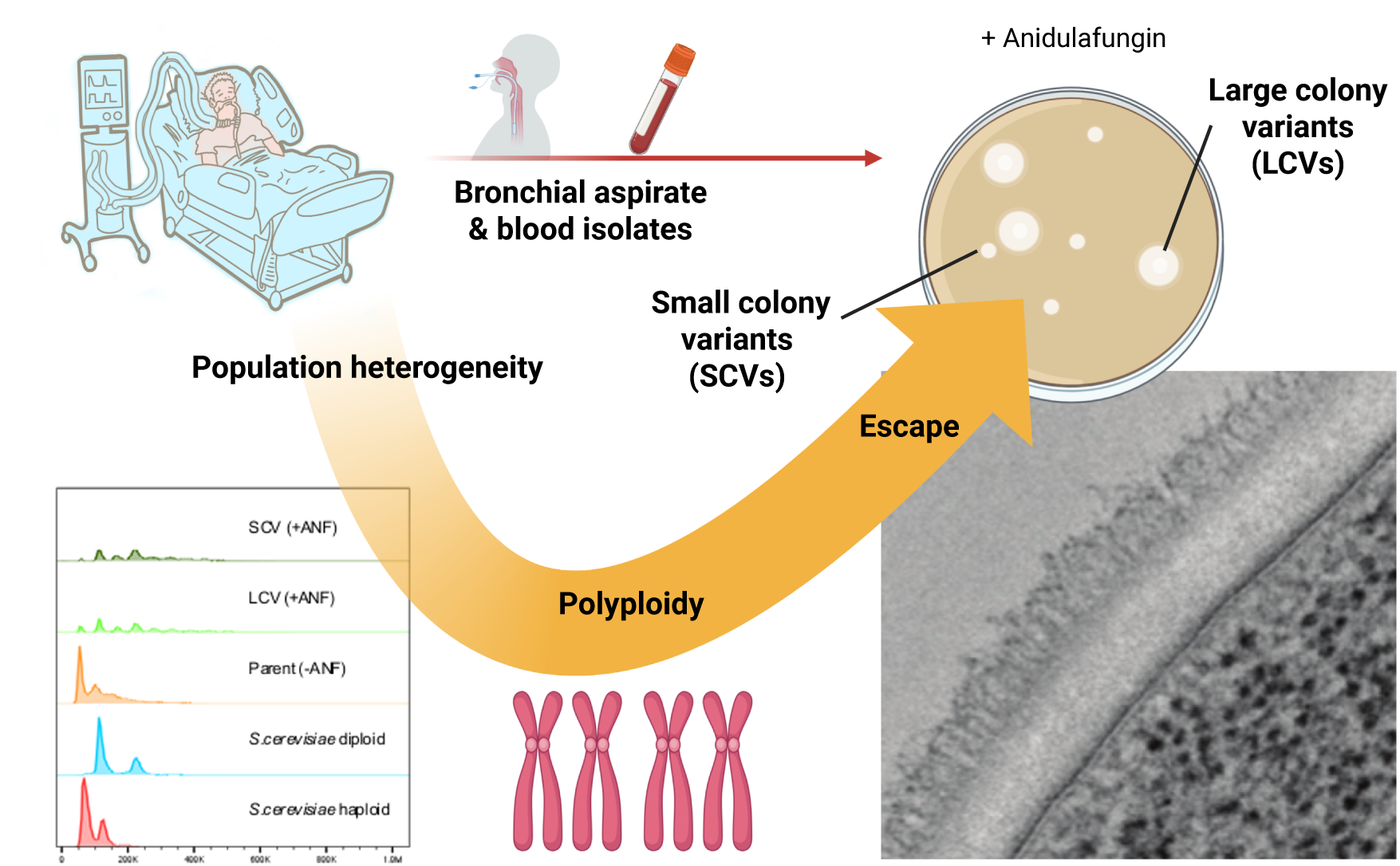

## Introduction

*Candida auris*, also named *Candidozyma auris*^1^, is a World Health Organization critical priority human fungal pathogen^2^. This fungus causes invasive infections (candidiasis) with an associated mortality of up to 45%^3^, and propensity for resistance to all key antifungal classes: azoles, echinocandins, and polyenes^4,5^. After detection in Japan in 2008^6^, six clades have emerged near-simultaneously across the world^7,8^. *C. auris* colonises the skin, particularly of patients who are critically ill or in long-term ventilation facilities^9^. Furthermore, *C. auris* is challenging to eradicate from hospital environments, resulting in inter-patient spread within and between hospitals^10–13^. Colonisation and infection rates continue to rise (in Europe^14^, North America^15^, South America^16^, Africa^17^, and Asia^18^) with a recent resurgence in UK cases prompting the inclusion of *C. auris* in the list of notifiable organisms, updates to guidelines for healthcare settings, and the declaration of a national public health incident in 2025^19,20^.

As the majority of *C. auris* isolates are resistant to fluconazole (>97% for clades I and III) and a high proportion to amphotericin B (47% for clade I)^21^, echinocandin monotherapy is currently the initial treatment recommended for invasive infections^5,22,23^. Initially, echinocandin resistance was uncommon across clades I-IV (0-9%)^21^, but cases continue to emerge. Echinocandins target the essential cell wall enzyme β-glucan synthase, encoded by *FKS1* and *FKS2* in *C. auris*^24^. Resistance is normally defined as growth above a minimum inhibitory concentration (MIC) using epidemiologically-derived or clinical breakpoints based on correlation with clinical outcomes^25,26^. Typical resistance is thought to arise primarily through mutations in *FKS1* hot-spots^27–31^ that lead to treatment failure *in vivo*, as demonstrated in murine infection models^32,33^. Several clinical case studies have shown resistance arising due to *FKS1* mutations: in a five isolate series from a single patient over 1 year, only the terminal isolate contained the F635Y *FKS1* variant and was echinocandin resistant^34^. In a nineteen isolate series from a single patient over 72 days, F639Y/F635C variants emerged alongside resistance to four classes of antifungal (pan-resistance) with further mutations in genes related to azole, polyene, and flucytosine resistance^35^.

Echinocandin resistance can also be caused by *FKS1/2*-independent mechanisms that either promote resistance directly, or that lead to *FKS* mutations. In four cases of urinary infection, *FKS1* mutation was necessary for echinocandin resistance, but other mutations related to cell wall stress and DNA repair/chromatin remodeling were present in strains with elevated MICs^36^. *In vitro* micro-evolution experiments have shown additional mutations outside *FKS1* hot-spots, including in *ERG3*, associated with echinocandin resistance^37,38^. Large scale genome-wide association studies have suggested that mutations in other cell wall-related genes (*IFF4*, *FCR1* and *GWT1*) may promote echinocandin resistance^39^. In *Candida albicans*^40^ (and other fungal pathogens^41,42^), aneuploidy and copy number variation can also lead to echinocandin resistance. Such structural variation has not yet been described in *C. auris* with respect to echinocandin resistance^43^. Furthermore, the development of secondary echinocandin resistance, and even pan-resistance to azoles, polyenes, echinocandins and flucytosine, has been reported in patients receiving treatment for clinically refractory infection, described in as many as 3% of clade I isolates^15,21,35,44,45^. The step-wise mechanisms underlying the emergence of echinocandin resistance remain poorly understood and are urgent questions for *C. auris* therapy in light of such limited options^43,46^.

In addition to standard antifungal drug resistance, heteroresistance and tolerance phenomena can contribute to population heterogeneity, stress adaptation, and treatment failure in infections caused by other pathogenic *Candida* species^47^, including *C. parapsilosis*^48^, *C. glabrata*^49^ and *C. albicans*^50^. Heteroresistance describes a small intrinsically resistant sub-population of cells (<1%) that is selected for and expands under drug pressure^47,48,51^. Tolerance represents the ability of a larger sub-population of cells (10-50%) within an isogenic, drug susceptible population (using MIC) to survive and grow slowly (>24 h) at concentrations above the MIC^47^. In other pathogenic *Candida* species, echinocandin tolerance has been shown to arise *via* increased chitin synthesis and cell wall remodeling triggered by protein kinase C (PKC), Hog1, Hsp90 and calcineurin signaling^52,53^. In *C. auris*, echinocandin exposure *in vitro* can lead to the increased expression of genes involved in cell wall synthesis and remodeling including genes encoding chitin synthases, cell wall adhesin Als5 and the drug efflux pump Cdr1^24,54,55^. These transcriptional changes correlate with echinocandin-induced adaptations such as cellular adhesion, aggregation, and biofilm formation^54–56^.

Here, we explore underlying mechanisms associated with the emergence of echinocandin refractory infection in two critically ill patients with *C. auris* bloodstream infections. Our data provide evidence for the sequential accumulation of a series of physiological and genetic adaptations leading to the emergence of resistance and echinocandin treatment failure.

## Results

### *FKS1* mutations and aneuploidy are insufficient to account for echinocandin refractory *C. auris* infection

We identified *C. auris* isolates from two patients with bloodstream infections (BSI) refractory to echinocandin therapy treated in the intensive care units of St George’s (isolates StG1-5, **Figure 1A**), and King’s College Hospitals (isolates K1-2, **Figure 1B**) in London, UK. The first patient suffered persistent candidaemia for three successive days on treatment, whilst the second patient experienced an initial BSI, followed by breakthrough infection on echinocandin treatment after 33 days. Antifungal susceptibility testing performed by the UK Mycology Reference Lab in Bristol suggested that these fungal isolates were not echinocandin resistant (**Figure 1C**) based on the tentative CLSI breakpoint (≥4 μg/mL) for anidulafungin in *C. auris*^57^. However, anidulafungin monotherapy failed to clear the fungal infection and both patients required a switch to amphotericin B-based combination treatment and subsequently died. Therefore, we sequenced the genomes of these *C. auris* isolates to explore the basis of their lack of response to echinocandin therapy.

**Figure 1:**
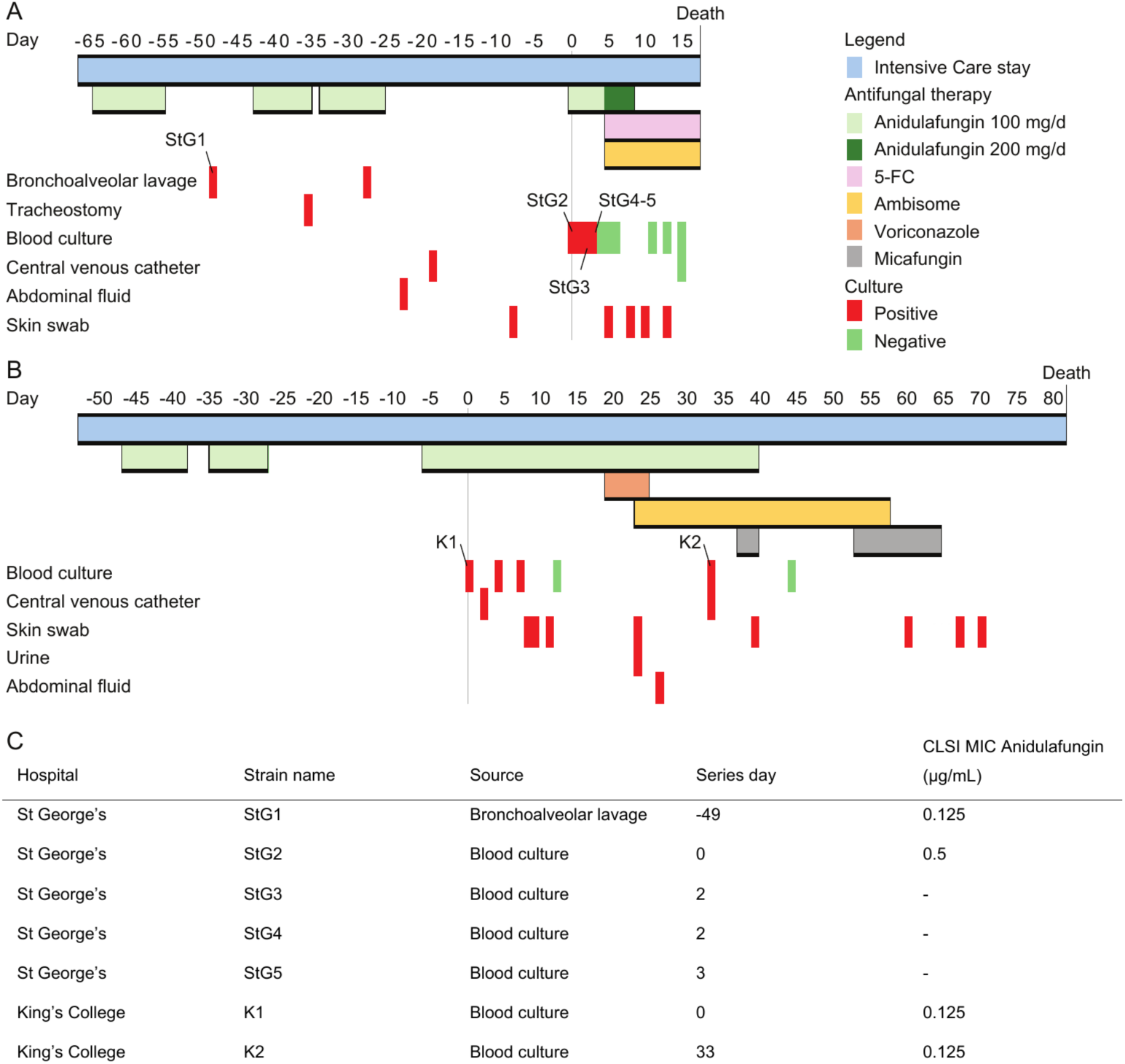
Clinical case series for patients suffering from *C. auris* candidaemia during antifungal therapy, both pre- and post-clinical isolate collection. **(A)** St George’s Hospital case series (isolates StG1-5), beginning at day -67 from first blood culture isolation (day 0), and ending at death on day 17. **(B)** King’s College Hospital case series (isolates K1-2), beginning at day -53 before first blood culture and ending at their death on day 81. This case was previously included (case no. 10) in an outbreak report^11^. **(C)** Clinical isolates included in this study by hospital site, day of isolation and CLSI MIC values against anidulafungin as reported by NHS Microbiology laboratories.

To understand the genetic mechanisms underpinning echinocandin therapy failure in these two patients, we sequenced the genomes of their *C. auris* isolates. Whole genome sequencing revealed that each of the isolates belonged to clade I (**Figure S1**) and, surprisingly, no non-synonymous mutations were present in *FKS1/2* genes. Therefore, we sought to understand additional pathogen-related mechanisms by which these infections were refractory to treatment. We identified 2,567 single nucleotide polymorphisms (SNPs) compared to the B8441 clade I reference genome, including 2,104 in both StG and K series, of which 198 were present only in StG isolates, and 264 only in K isolates. Aside from SNPs in intergenic regions (71.0%), there were a total of 469 non-synonymous variants, 6 nonsense variants, and 66 indels in coding regions (**Figure 2A-B**). The greatest variation across clinical isolate series was observed in *HYR3* (B9J08_004100: one deletion, four non-synonymous variants, and nine synonymous variants), which is predicted to encode a GPI-anchored cell wall protein^58^ (**Figure 2A-B**). We also identified mutations in genes associated with cell wall stress response (*BCK1*), cell wall synthesis (*KAR2*, *RBR3*, *SCF1*), secretory pathways and intracellular trafficking (*GDI1*, *SVL3*, *VPS5*, *YPT6*), RNA synthesis (*HAS1*, *IWR1*), and histone acetylation (*SPT10*), suggesting genetic adaptations involving cell wall remodeling in response to drug pressure (**Table S1**). SNPs resulting in non-synonymous mutations were identified in genes related to azole resistance: *ERG11* (K143R), *CDR1* (V704L) and *TAC1b* (A640V) in isolates StG1-5; and *ERG11* (Y132F), *CDR1* (E709D) and *TAC1b* (A583S) in isolates K1-2. No non-synonymous mutations were observed in *MDR1-2*, *MRR1a-b*, *CAS5*, or *FUR1*.

**Figure 2:**
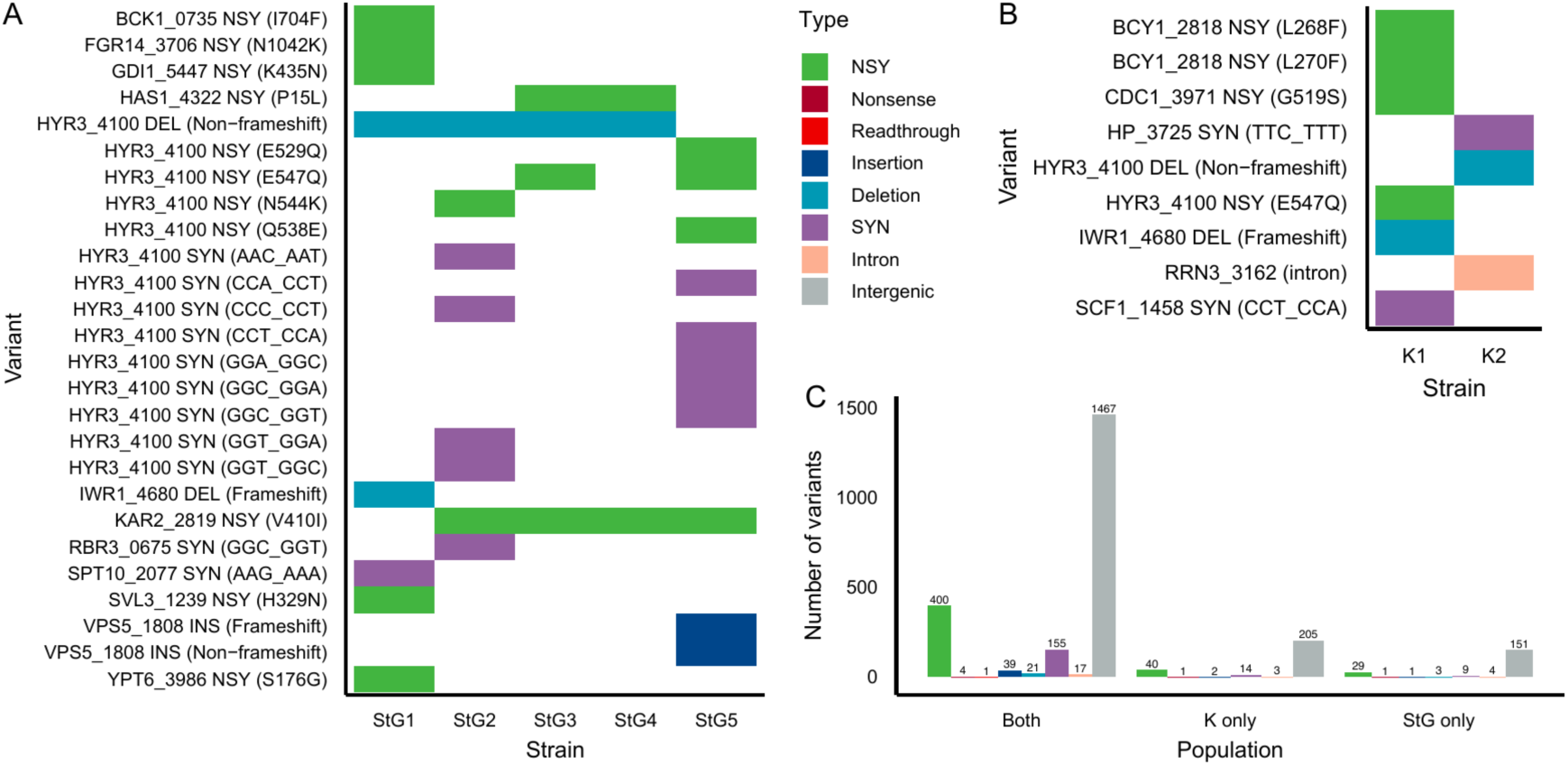
Common sequence variants identified in the genomes of the clinical series. **(A)** Genomic variants between clinical isolates in the StG1-5 series, excluding intergenic variants. **(B)** Genomic variants between clinical isolates in the K1-2 isolate series, excluding intergenic variants. **(C)** Counts of all genomic variants present across both clinical series.

We examined the genomes of the StG and K isolates (using thresholds of >1.4 and <0.6 normalised depth of coverage, **Figure S2A-B**) to identify gene or chromosome copy number variation (CNV) associated with echinocandin refractory infection. Though we did not identify aneuploidy, small regions showing increased per-gene CNV included loci such as *NTO1* (part of putative histone acetyltransferase machinery) and *FGR14* (homologue of a retroviral endonuclease-reverse transcriptase). Areas of low copy number include multiple genes (*e.g. CDC13*, *ECM21*, *ECM42*, *HEM14*, *IWR1*, *MRF1*, *MRP49*, *PRD1*, *RCO1*, *TMA17*). To our knowledge, none of these CNV-affected genes have been associated with echinocandin resistance (**Table S2**).

### *C. auris* isolates display echinocandin heteroresistance and rapidly develop further resistance *in vitro*

Based on CLSI breakpoints, both StG and K isolates were echinocandin sensitive and yet obtained from clinically refractory infections. Therefore, we tested whether initial StG1 and K1 isolates displayed anidulafungin heteroresistance using population analysis profiling (PAP, **Figure 3A**). Heteroresistance was indicated by the growth of small sub-populations of cells (<0.01%) on YPD-agar at anidulafungin concentrations 256-fold greater than those for the susceptible majority (**Figure 3A**). After extended culture (144 h), two distinct colonial morphologies emerged that were most pronounced on YPD-agar containing 8 μg/ml anidulafungin (**Figure 3B**): slower growing “small” colony variants (SCVs), and faster growing “large” colony variants (LCVs).

**Figure 3:**
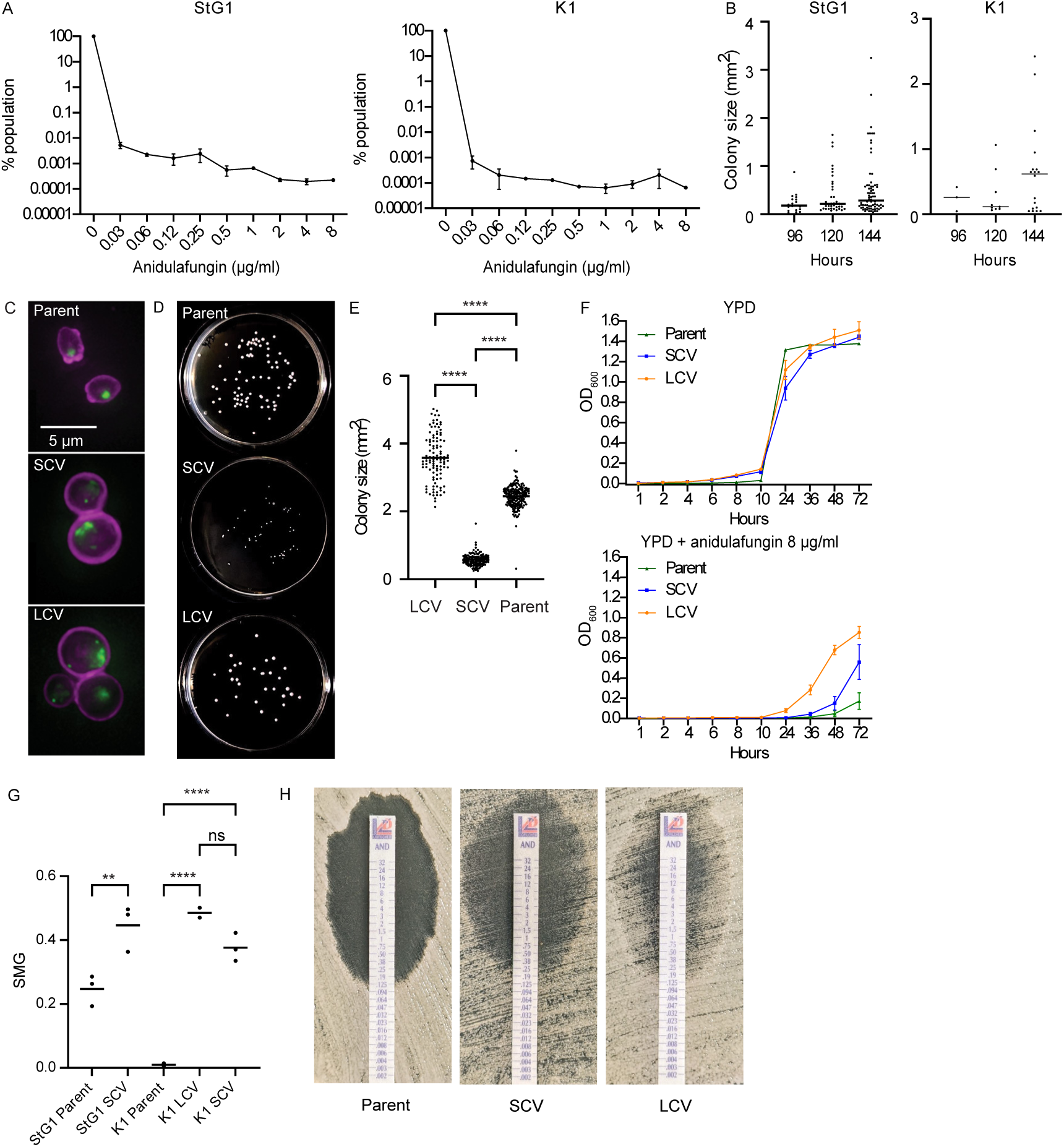
*C. auris* isolates display heteroresistance with sub-populations of cells displaying contrasting colonial morphotypes. **(A)** Population analysis profiling (PAP) of StG1 and K1 clinical isolates demonstrates anidulafungin heteroresistance. Points represent mean of 3 biological replicates. Error bars represent standard error of the mean (SEM). **(B)** Emergence of small and large morphotypes after protracted growth under drug pressure. StG1 and K1 isolates were plated onto YPD containing 8 μg/mL anidulafungin, and colony sizes measured using ImageJ after 96, 120 and 144 h incubation (biological triplicates). **(C)** Fluorescence microscopy of individual cells, their cell walls stained with CFW and their DNA content with SYBR green. **(D)** Colony morphotypes of StG1 are stable following re-plating. StG1 large colony variants (LCVs) and small colony variants (SCVs) were taken from a YPD+anidulafungin 8 μg/mL plate, and the StG1 parental isolate taken from a YPD plate. These cells were replated onto YPD at 10^2^ CFU and imaged after 48 h at 30 °C. **(E)** Colony sizes were then measured on these plates (**F**) by ImageJ. Plots represent distribution of all colony sizes from 3 biological replicates for each cell type. One-way ANOVA: *, *p* <0.05; **, *p* ≤0.01; ***, *p* ≤0.001; ****, *p* ≤0.0001. **(F)** Growth (OD_600_) of the StG1 parent, LCV and SCV colonies in YPD containing 0 or 8 μg/ml anidulafungin. Points represent mean of 3 biological replicates, error bars represent SEM. **(G)** Supra-MIC growth for StG1 SCVs *vs* parent isolate. Each point represents mean of technical triplicate results. One-way ANOVA, *p*-values as above. **(H)** Etest strips reveal the zones of inhibition for the parental isolate StG1, LCVs and SCVs.

We compared the phenotypes of SCVs and LCVs: fluorescence microscopy revealed that, in the presence of anidulafungin, both SCVs and LCVs contained moderately larger cells than the parental isolate alongside aggregation and DNA content increase on SYBR green staining (**Figure 3C**). The SCV and LCV morphotypes persisted when they were re-plated onto drug-free plates: re-plated SCVs (mean colony size 0.58 mm^3^) formed significantly smaller colonies (*p* < 0.0001) than LCVs (mean colony size 3.58 mm^3^) and parent isolates (mean size 2.45 mm^3^) following 48 h growth at 37 °C (**Figure 3D-E**). SCVs and LCVs grew at similar rates to parental isolates in drug-free liquid YPD. However, in YPD containing 8 μg/mL anidulafungin, SCV cells grew significantly faster than parental cells, and LCVs even faster than SCV cells (**Figure 3F**). Furthermore, both SCVs and LCVs were resistant to anidulafungin according to updated EUCAST criteria for *C. auris* (sensitive, S: ≤ 0.25 μg/mL; resistant, R: >0.25 g/mL^25,26^): MIC against anidulafungin was highest for StG1 and K1 LCVs (modal MIC 4-8 μg/ml, range 2-16 μg/ml); followed by SCVs (modal MIC 2 μg/mL, range 1-4 μg/mL) and parental isolates (modal MIC 1-2 μg/mL, range 1-4 μg/mL, **Table S3**). SCVs also exhibited significantly higher tolerance to anidulafungin compared to the parent isolate, measured using both supra-MIC growth (**Figure 3G**) and confirmatory Etest inhibition strips (**Figure 3H**). LCVs were resistant according to Etest (**Figure 3H**). According to the updated EUCAST guidelines, parental isolates were also resistant.

Taken together, these findings indicate the presence of stable heterogeneous sub-populations within treatment-refractory *C. auris* clinical isolates, which are consistently isolated at high anidulafungin concentrations. These distinctive small and large colonial morphotypes appear to reflect a spectrum of drug-adapted sub-populations, as shown by increased growth rates in the presence of drug, higher tolerance and anidulafungin MICs-compared to the parent (rather than a difference in actual cell size).

### *CRZ1* mutations in LCVs are associated with anidulafungin resistance

We explored the genomic foundations of the colony morphotypes with associated tolerance and resistance phenotypes by sequencing SCV (*n* = 18) and LCV (*n* = 17) colonies. Mutations potentially underlying the observed population heterogeneity were identified by comparing their genomes with those of parental controls: StG1, K1 and K2 grown on YPD, with additional SCV/LCV isolates sequenced directly on anidulafungin at 8 μg/ml (*n* = 3 per isolate). No *FKS1/2* mutations were observed in any of the SCV or LCV genomes, indicating that anidulafungin resistance had emerged *via FKS1/2*-independent mechanisms.

To seek these *FKS1/2*-independent mechanisms, we examined sequence variants displaying a frequency difference of >25% between groups (*e.g.* SCV *vs* LCV, **Figure S3A**) and compared the number of strains in each group that displayed variants (**Figure S3B-C**, *Supplementary Note 1*). Strikingly, variation in the *CRZ1* gene was significantly more common in LCVs compared to parent isolates (14/17 *vs* 0/7, 82.3% *vs* 0%, adjusted *p*-value = 0.032) and between LCVs and SCVs (2/18, 11.1%, adjusted *p*-value = 5.93 x10^-5^). A total of 16 non-synonymous mutations were identified in *CRZ1*, excluding S237Y which was present in all strains (**Figure 4**). The majority of *CRZ1* mutations were highly likely to alter function, including nonsense (*n* = 8) and frameshift deletions (*n* = 3), compared to non-synonymous mutations (*n* = 5). Thus LCVs, which were anidulafungin resistant, carried the largest number of *CRZ1* mutations (**Figure 3F**), consistent with the recent finding that *C. auris crz1* knockout mutants are anidulafungin resistant^59^.

**Figure 4:**
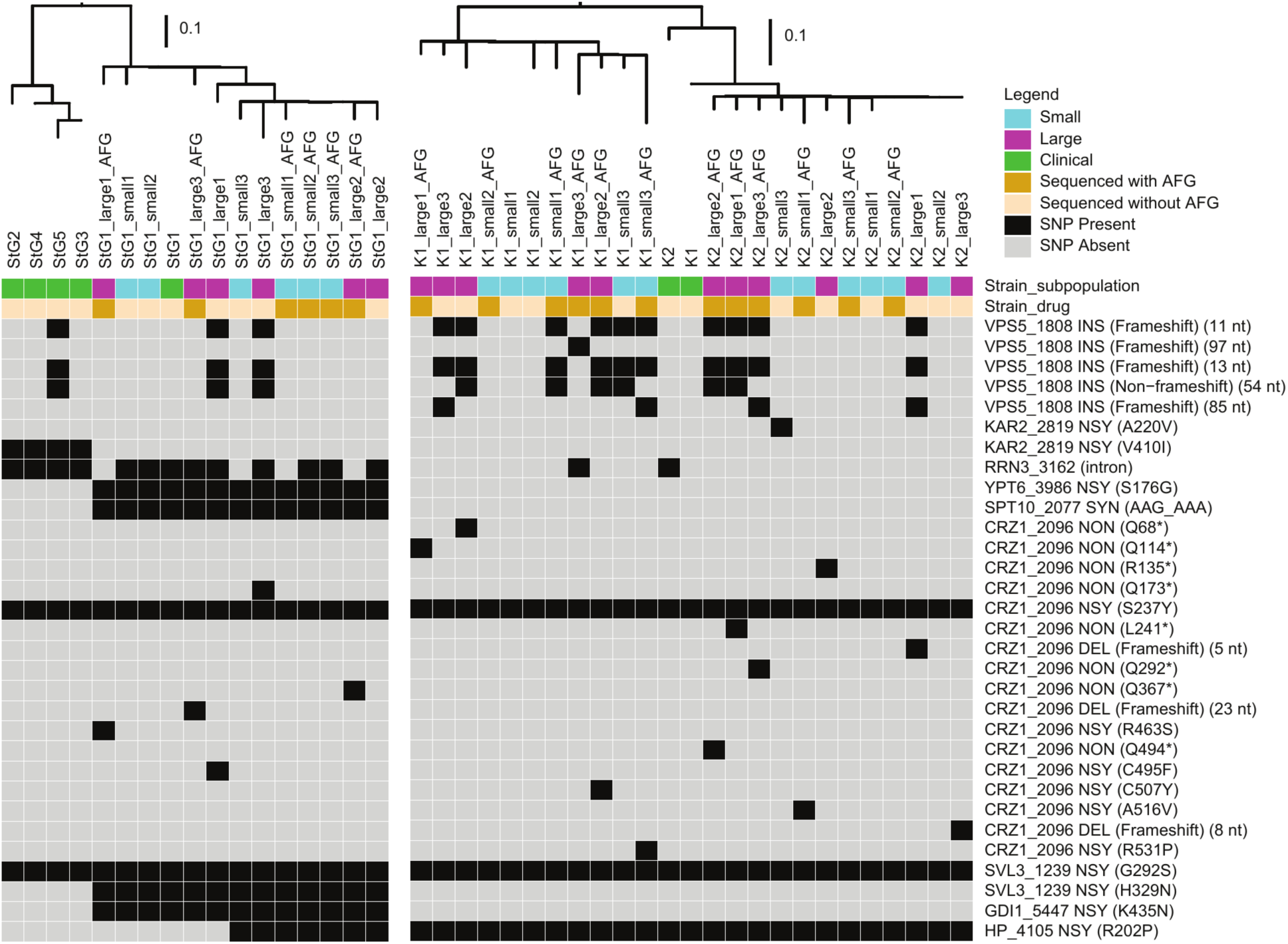
Significant within-gene variation in clinical isolates and their large and small morphotypes. FastTree phylogenies for StG and K clinical series isolates are shown above. The colours in the legend (top right) and below the phylogenies indicate the group: clinical isolate versus large or small colony variant; genome sequenced following growth overnight on- or off-anidulafungin (8 μg/mL). Black boxes indicate the presence of the variant specified to the right of the figure: NSY: non-synonymous, SYN: synonymous, NON: nonsense, INS: insertion, DEL: deletion, AFG: anidulafungin. Differences between isolates K1 and K2 were driven by 237 intergenic variant differences (95 present only in isolate K1, 142 present only in isolate K2) in addition to those outlined in Figure 2B.

The *CRZ1* calcineurin-responsive transcription factor gene was one of two genes in the calcineurin pathway that displayed highly significant variation in SCVs and LCVs. Another calcineurin-pathway related gene (*RCN2*, regulator of calcineurin) was significantly enriched for associated intergenic SNPs that flanked this gene in LCVs (88.2%) *vs* SCVs (27.8%, adjusted *p*-value 0.017). Furthermore, four frameshift mutations in *VPS5* (predicted vacuolar sorting protein) were identified in fourteen strains, but these were not significantly enriched in SCVs or LCVs relative to the parental isolates. However, two non-synonymous variants were significantly enriched in daughters compared to clinical parents: *KAR2* V410I (*p* = 0.0077) and *HP_4105* R202P (*p* = 0.033). The change in Kar2 (a member of the Hsp70 chaperone family) could conceivably contribute to the tolerance profile of *C. auris* through stress response adaptations. However, the function of *HP_4105*, and hence the potential impact of the R202P mutation, remains obscure.

### Anidulafungin induces cell wall remodeling in both SCVs and LCVs

In *C. albicans*, the inhibition of β-1,3-glucan synthesis by echinocandins induces compensatory increases in chitin synthesis and cell wall remodeling^60^, in part *via* Crz1 signaling^61^. In *C. auris*, caspofungin induces chitin synthase gene expression in a *CRZ1*-dependent fashion^59,62^. Therefore, we examined the impact of anidulafungin on the cell walls of LCVs and SCVs. Flow cytometry of Calcofluor-White (CFW) stained cells revealed that chitin levels increased in response to anidulafungin in both LCVs and SCVs (**Figure 5A**), with associated increases in β-1,3-glucan exposure (**Figure 5B**). Transmission electron microscopy (TEM) showed corresponding changes in cell wall architecture; exposure to anidulafungin led to significant thickening of the inner (chitin-rich) cell walls of both LCVs and SCVs (**Figure 5C-E**). Anidulafungin also induced a slight increase in the outer mannan layer of the cell wall, but this was minor compared to the dramatic changes to the inner layer. High pressure ion chromatography (HPIC) of cell wall carbohydrates provided further evidence of anidulafungin-induced increases in chitin/mannan content with corresponding decreases in glucan both across both the StG clinical isolates (**Figure 5F-H**) and in SCVs, but not in LCVs (**Figure 5I-K**).

**Figure 5:**
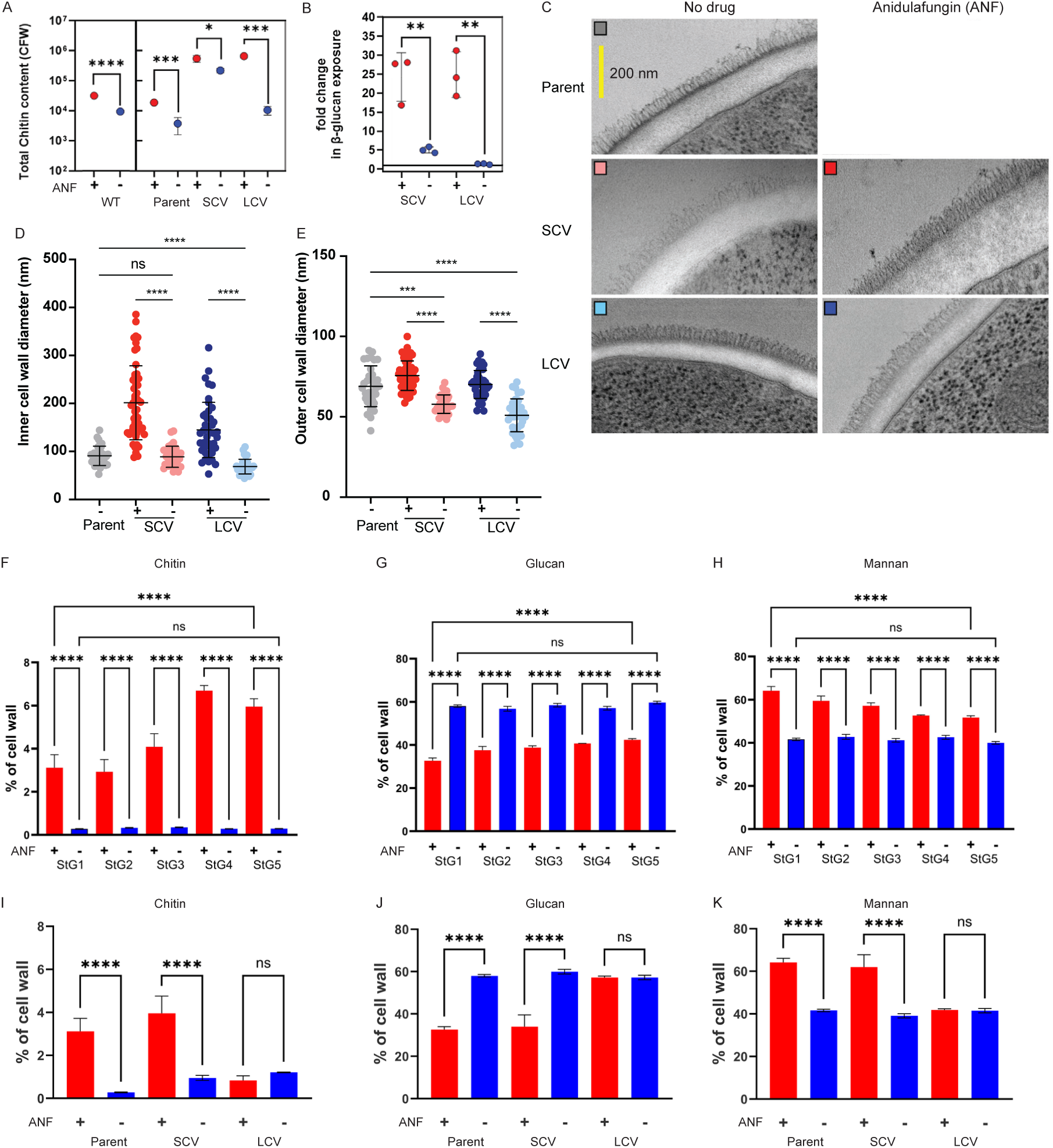
Impact of anidulafungin on chitin content and β-1,3-glucan exposure of cell walls of StG parents and large and small morphotypes: **(A)** The chitin content of the cell wall of StG parents and large and small morphotypes during growth in the presence (+, red) or absence (-, dark blue) of 8 μg/ml anidulafungin (ANF) was quantified by Calcofluor-White (CFW) staining and flow cytometry. Data represent means and standard deviations from three replicate experiments, one-way ANOVA: ns, not significant; *, *p* <0.05; **, *p* ≤0.01; ***, *p* ≤0.001; ****, *p* ≤0.0001. **(B)** The fold change in β-1,3-glucan exposure in large and small morphotypes, compared with parental isolates, in the presence (+, red) or absence (-, dark blue) of anidulafungin was quantified by dectin-1 staining and flow cytometry. Statistics as per **(A)**. **(C)** Transmission electron microscopy (TEM) reveals cell wall remodeling in response to anidulafungin. The diameters of the **(D)** inner and **(E)** outer cell wall layers were quantified from TEM images using ImageJ. Data represent means and standard deviations from *n* >30 cells (10 measurements per cell) and were analysed using Brown-Forsythe and Welch ANOVA, *p*-values as above. **(F-K)** High pressure ion chromatography (HPIC) was used to measure the chitin (**F**), glucan (**G**) and mannan (**H**) contents of the cell walls of the StG1-5 series following growth on YPD containing 0 or 8 μg/mL anidulafungin. The chitin (**I**), glucan (**J**) and mannan (**K**) contents of the cell walls of the parental isolate StG1 and its SCV and LCV isolates and their corresponding changes in cell wall content inc. chitin, glucan, and mannan after growth without or with (8 μg/mL) anidulafungin. One-way ANOVA, *p*-values as above.

### Clinical and drug-tolerant/drug-resistant sub-populations of *C. auris* are characterised by alterations in ploidy

Stress-induced changes in ploidy have been proposed to precede the emergence in drug resistance in *C. albicans*^63,64^, and tetraploidy has been described in *C. albicans* clinical isolates from human hosts^65–68^. Therefore, we compared the ploidies of *C. auris* StG1 and its SCV and LCV daughters, analysing cells taken directly from colonies on plates using flow cytometry: StG1 cells from YPD plates, and SCV and LCV cells from were taken directly from YPD plates containing 8 μg/ml anidulafungin. *Saccharomyces cerevisiae* haploid and diploid strains were used as controls. The *S. cerevisiae* controls showed clean 1n-2n and 2n-4n ploidies, as expected for mixed populations containing cells pre- and post-S-phase (**Figure 6A-B**). Although *C. auris* is purportedly haploid^69,70^, StG1 cells displayed heterogeneous ploidies ranging from 1n to >4n (**Figure 6A-B**). Even higher ploidy ranges, with a higher proportion of cells with ploidies of >4n, were observed for the SCV and LCV morphotypes isolates from YPD+anidulafungin. Our bioinformatic analyses, which included allele frequency tallies (**Figure S4A**), were consistent these increased ploidies, though identification of heterozygous sites was not sensitive to inter-strain differences (**Figure S4B**).

**Figure 6:**
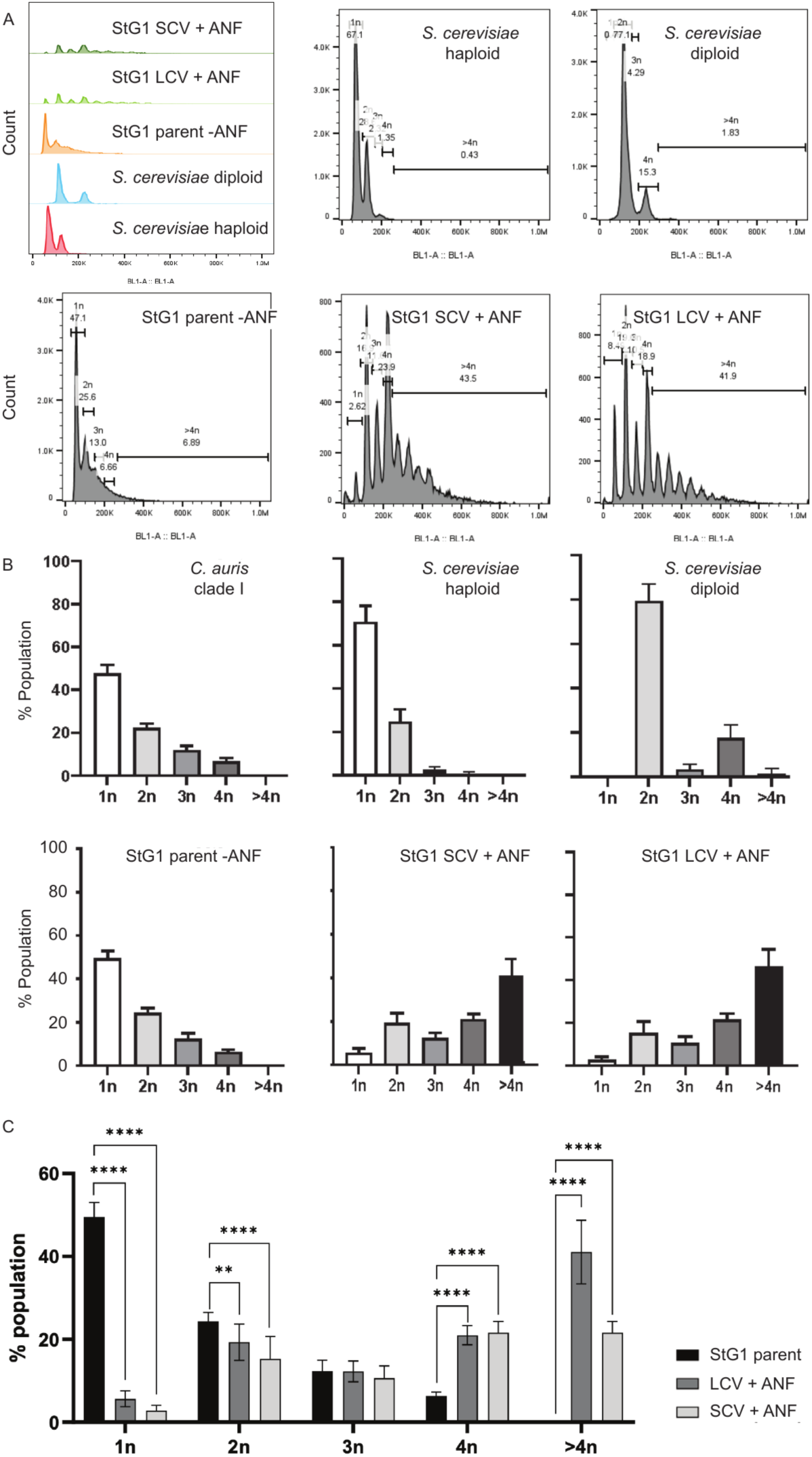
Exposure to anidulafungin increases the ploidy of *C. auris*. **(A)** The ploidy of *C. auris* cells in colonies taken directly from plates was measured by staining their DNA content with SYBR green and quantifying the fluorescence by flow cytometry. Haploid and diploid *S. cerevisiae* strains and a *C. auris* clade I isolate were used as controls. Colonies for the *C. auris* clade I and *S. cerevisiae* controls were taken from YPD plates. Colonies for the *C. auris* StG1 parent and its small and large colony variants were harvested from YPD plates containing 8 μg/mL anidulafungin. Peak volumes were quantified using were quantified using FlowJo v.10 software. (**B**) The proportions of 1, 2n, 3n, 4n and >4n cells in each sample were calculated from the peak volumes obtained by flow cytometry in (**A**) with unpaired t-test comparison between groups (**C**): *p* <0.05; **, *p* ≤0.01; ***, *p* ≤0.001; ****, *p* ≤0.0001.

The heterogeneous ploidy of StG1 cells was unexpected (**Figure 6A-B**). Therefore, to test whether other *C. auris* isolates display this phenotype, we examined isolates from clades I to V. Ten epidemiologically divergent isolates all displayed heterogeneous ploidies when grown on YPD agar (**Figure S4C**) with strict single-cell gating strategies (**Figure S4D**). These findings suggest that genomic instability and polyploidy are features of *C. auris* that might underlie this species’ phenotypic heterogeneity and propensity towards drug tolerance/resistance, promoting treatment-refractory infection and emergence of secondary echinocandin resistance.

## Discussion

In this study we investigated serial invasive *C. auris* isolates from critically ill patients, which did not respond to anidulafungin despite the absence of *FKS* mutations. We have identified the presence of phenotypically heterogeneous sub-populations in *C. auris*, which display distinct colony morphotypes that show increased ploidy and enhanced adaptation to drug stress, relative to the parent clinical isolate. The larger colony variant (LCV) grows faster in the presence of anidulafungin, has an elevated anidulafungin MIC, and carries significantly more *CRZ1* mutations than the small colony variant (SCV) and parental isolate. Whilst *C. albicans* and *C. glabrata crz1* mutants display echinocandin sensitivity^71–73^, a recent study has suggested a *C. auris crz1* null mutant is anidulafungin-resistant^59^. Consistent with this, *CRZ1* mutations observed in LCVs were nonsense/stop or deletion/frameshift mutations, which likely result in Crz1 disruption/dysfunction, and these were associated with increased anidulafungin resistance. *CRZ1* is likely to promote the fitness of *C. auris* under drug pressure, but *C. auris crz1* knockout strains do not display attenuated virulence in a murine model of systemic candidiasis^59^.

Based on our findings, we propose that the following mechanisms contributed to the failure of anidulafungin therapy in the patients in our study. A small sub-population of drug-tolerant *C. auris* was able to survive high supra-MIC concentrations of anidulafungin (giving rise to SCVs). This sub-population of cells was able to adapt to anidulafungin by elevating chitin synthesis and thickening their inner cell walls, which may have contributed to the drug tolerance/intermediate resistance phenotype of SCVs. We reason that the survival of these cells under drug pressure then enabled the emergence of genetic resistance, in part *via CRZ1* escape mutations, to yield the faster growing, anidulafungin resistant LCVs. The Crz1 transcription factor regulates cell wall genes in many fungi^73^, and therefore the loss of Crz1 functionality likely contributed to the observed blunting of the cell wall remodeling response of LCVs to anidulafungin.

Previous reports have suggested that echinocandin resistance in *C. auris* primarily arises *via* mutations in *FKS1/2* hot-spots *in vivo*^27–31^. We did not identify *FKS1/2* mutations in clinical isolates or sub-populations of SCVs/LCVs. Instead, we observed a predominance of *HYR3* variation within the clinically evolved case series. *HYR3* encodes a predicted GPI-anchored cell wall protein that is under selection in BSI-causing *C. auris* clades I, III and IV^58^. Also, *HYR3* was highly upregulated in a murine catheter infection biofilm model *in vivo*^74^, and in mature biofilms with coincident echinocandin resistance^75^. Our findings reinforce the view that this locus is significant during human infection.

Variable ploidy is likely to be an important aspect of generating phenotypic heterogeneity in *C. auris*, and is perhaps an overlooked feature of genomic plasticity in this pathogen. Reports of polyploidy in *C. auris* are rare^70^, though diploid isolates have been reported from clades I and III, which are associated with higher virulence in a murine systemic infection model^76^. Polyploidy can accelerate genomic evolution and is viewed as a common and reversible fungal stress response that can increase the availability of beneficial mutations^41,42,65,77–79^ and even potentially increase virulence^63,80^. We provide evidence for drug-induced increases in ploidy in treatment-refractory *C. auris* isolates, strongly suggesting that changes in ploidy promote therapeutic escape by this important fungal pathogen^69,70^.

In light of our findings, we suggest updates to the three-stage model for the development of echinocandin resistance in *Candida* species^36,81^. We propose that the emergence of resistance begins with echinocandin tolerance driven by reversible physiological changes, for example *via* protein kinase C, calcineurin-Crz1, HOG and Hsp90 signalling^82^. Then, the exposure of sub-populations of physiologically relatively tolerant cells to sub-MIC echinocandin concentrations, especially in difficult-to-penetrate sites (*e.g.* catheter-related biofilms or intra-abdominal compartment, as was the case for our two critically ill patients), selects for the progressive emergence of resistance. Here, drug-induced increases in *C. auris* ploidy promote the accumulation of escape mutations. Additional drug pressure would then select for further mutations, for example in *FKS1* and/or *CRZ1*, that promote increased resistance and faster growth, thereby yielding the LCV phenotype. This updated model can account for the observed population heterogeneity of clinical isolates from patients that have undergone protracted drug treatments, where parental cells coexist with SCVs and LCVs displaying varying degrees of tolerance and resistance and may represent parallel pathways towards overcoming antifungal pressure. This update may also inform the developing conceptualisation of bet-hedging in fungal pathogens^83^.

Our investigation of fungal population heterogeneity in patient isolates has revealed progressive pathways towards the emergence of echinocandin resistance in *C. auris* that may otherwise have been overlooked. The causes of treatment failure are complex, involving also drug delivery to the site of infection, as well as host immunosuppression associated with critical illness. Nevertheless, defining the mechanisms that underlie the emergence of drug tolerance and resistance in fungal pathogens, together with the temporal dynamics of their impacts *in vivo*, may ultimately lead to improved patient outcomes. Therapeutic strategies could be tuned in real time, to minimise the development of drug tolerance or resistance in an individual patient^47,84^. Combination therapy is one such powerful option that could enhance efficacy, addresses intrinsic fungal population heterogeneity, and reduce the inherent risk of resistance selection by currently recommended echinocandin monotherapy^85,86^: for example, the ongoing Wellcome-Trust funded COMBAT *Candida* clinical trial will compare micafungin alone to combination with micafungin and flucytosine for treatment of candidaemia in a setting of high *C. auris* prevalence in South Africa, incorporating resistance emergence as an endpoint.

In summary, we have uncovered pathogen-based mechanisms whereby *C. auris* sub-populations adapt to first-line therapy, contributing to persistence and evolution. The identification and association of *CRZ1* mutation with resistance provides a valuable candidate locus for future functional investigation, potentially serving as an early marker of resistance evolution. Future research should focus on validating *CRZ1* mutations by using structural biology, gene editing, and population genetics approaches. The clinical implications of our findings are that treatment refractory *C. auris* infections require a more nuanced approach to resistance beyond MICs, and novel therapeutic approaches are needed to address the array of adaptive mechanisms that *C. auris* displays in the face of echinocandin pressure.

## Methods

### DNA Extraction

Clinical isolates were confirmed as *C. auris* using MALDI-TOF and tested for MIC by CLSI standards, stored at -80 °C in YPD containing 20% glycerol, and sub-cultured at 30 °C on YPD agar (Sigma Aldrich, UK). Yeasts were grown overnight in 10 ml YPD at 30 °C before centrifugation at 13,000 rpm, resuspension in 3 ml water and transferring into a 2 ml tube. Cells were centrifuged at 13,000 rpm for 5 min. prior to the supernatant being removed and cells resuspended in the residual liquid. Cells were snap-frozen by immersion in liquid nitrogen for 1 min. and then transferred into a 65 °C water bath for 3 min. Disruption buffer included 400 mg of glass beads, 200 μl of DNA extraction buffer (2% Triton X-100, 1% SDS, 100 mM NaCl, 1 mM EDTA, 10 mM Tris HCl (pH 8)), and 200 μl of phenol:chloroform:isoamyl alcohol (25:24:1), which was added to yeast samples, which were vortexed for four rounds of 20 s with 1 min. breaks on ice to cool samples. A sample of 200 μl 1x TE was added to the mixture before vortexing again for a few seconds and centrifugation at 13,000 rpm for 10 min. The aqueous layer was transferred into a fresh tube and mixed with 1 ml of 100% ethanol and centrifuged at 13,000 rpm for 5 min. The pellet was resuspended in 0.4 ml TE and 3 μl of 10 mg/ml RNAase A and incubated for 15 min. at 37 °C. Samples of 10 μl of 4 mM ammonium acetate and 1 ml of 100% chilled ethanol were added. The mixture was centrifuged at 13,000 rpm for 10 min. at 4 °C. The supernatant was discarded, and pellet resuspended in 100% chilled ethanol. The mixture was spun at 13,000 rpm at 4 °C for 10 min. The supernatant was discarded, and the pellet was dried in a heat block at 65 °C for 10 min. DNA was resuspended in 50 μl water. DNA concentration was checked using Nanodrop and DNA quality by gel electrophoresis.

### Sequencing and variant calling

Sequencing was performed by the University of Exeter Sequencing Service (ESS) in the on the NovaSeq 6000 with SP flow cell (Illumina, San Diego, USA). Quality control was performed with MultiQC v1.10.1^87^. Variant Calling was performed using the Genome Analysis Toolkit (GATK) v4.1.2.0^88^ with alignment of raw sequences to the B8441 v2 reference genome (GCA_002759435.2)^7^ and the mitochondrial genome (NC_053321.1)^89^ using BWA-MEM v0.7.17^90^. HaplotypeCaller was executed in GVCF mode with the haploid ploidy flag. Hard filters were used to remove spurious variants, including the filters quality by depth (QD) <2.0, Fisher Strand (FS) >60.0 and root mean square mapping quality (MQ) <40.0.

### Population analysis profiling (PAP)

Overnight cultures of *C. auris* isolates were serially diluted 10-fold 6 times in sterile distilled water. 5 μl aliquots of each serial dilution were spotted in triplicate onto YPD agar plates infused with range of anidulafungin concentrations (0-8 μg/ml). Plates were incubated at 30 °C for 24-48 h and CFU counting performed to calculate CFU/ml able to grow on each drug concentration, which was then normalised to CFU/ml on drug free plate to calculate proportion of the population able to grow at each anidulafungin concentration. Due to noticeable heterogeneity in colony size at the maximum anidulafungin concentration (8 μg/ml), extended incubation of individual 8 μg/ml PAP plates inoculated with 10^7^ cells was performed. Plates were incubated at 30 °C for 7 days, and images taken after 48 h, 96 h and 144 h. Colony sizes were measured manually using ImageJ at each timepoint. Briefly, images were converted to black and white, sharpened, and then threshold changed to remove parts of the plate with no colony growing. To assess stability of SCV/LCV phenotypes, replica plating of isolates was performed by preparing inoculum of SCV/LCV in distilled water and then plating 10^2^ cells on drug free and 8 μg/ml anidulafungin-containing YPD agar. Plates were incubated at 30 °C for 48 h and colony sizes were calculated again using ImageJ as described previously.

### Growth curves

Microdilution plates were set up to contain 0, 0.25, or 8 μg/ml anidulafungin. Wells were inoculated with 1x10^5^ log phase cells such that the final volume was 200 μL/ well. Plates were incubated at 30 °C for 50 h and constantly shaken at 100 rpm. OD_600_ was measured using a Spectrostar Nano every 10 min. for 50 h. Prior to OD_600_ reading, the plates were shaken at 600 rpm.

### Susceptibility and tolerance assays

Clinical isolates were tested at the point of isolation using CLSI methodology confirmed at St. George’s and King’s College Hospitals with confirmation by the National Reference Laboratory in Bristol. Additionally, the susceptibility of isolates to anidulafungin was determined per EUCAST guidelines with slight modifications. Briefly, 96 well plates were prepared to contain two-fold dilutions of anidulafungin (Gereon B1224) with the first and last well containing no drug. Isolates were defrosted onto YPD agar and grown at 37 °C for 24 h. Five colonies were selected and resuspended in PBS and diluted such that 10^5^ cells were added to each well in a total volume of 200 μL/well except the last. Susceptibility in RPMI 1640 media supplemented with 2% dextrose was tested at 37 °C. OD_450_ was read using a Tecan plate reader. The tentative clinical breakpoints were used to determine whether an isolate was susceptible or resistant. Plates were also read at 48 and 72 h to determine supra-MIC growth (SMG). SMG was determined as average 72 h growth in the wells above MIC divided by the growth in the no-drug well. For Etest strips, isolates were grown for 48 h on YPD plates at 37 °C. Three to four colonies were taken and used to prepare 0.5 Mcfarland suspension. Sterile cotton swabs were used to streak each suspension across YPD plates in 3 separate directions to cover the whole plate in cells, and plates were allowed to dry. Sterile forceps were then used to place an Etest strip onto each plate, ensuring no air bubbles were present. Inoculated plates were incubated at 37 °C and were read and photos taken at 24 h and 48 h. Readings were made according to CDC guidelines for interpretation of Etest antifungal susceptibility testing results^91^.

### Phylogenomics

FastTree v2.1.11 with Jukes-Cantor modelling^92^ was used to construct phylogenetic trees based on multiple sequence FASTA alignments produced by ECATools^93^ using default parameters; midpoint rooted trees used 1,015 (StG clinical series plus laboratory evolved daughters) or 1,053 (K clinical series plus laboratory evolved daughters) phylogenetically informative sites. Additional sequences used to contextualise clinical samples within clades were obtained from the largest global genomic epidemiology study to date^21^ and the first five isolates were called and used for phylogenetic reconstruction, for which 192,032 sites were entirely covered in all.

### Statistical genomics and annotation

Copy number variation estimation and depth-of-coverage plots were calculated in comparison to normalised depth of coverage across the whole genome compared to the depth of coverage across either all positions for each gene locus or each 10 kb sliding window, from pileups created with Samtools mpileup^94^. Significance testing for variants/loci/functional annotations was performed using Fisher’s exact test (two-sided) with Benjamini-Hochberg (BH) testing for multiple correction with a cut-off of 0.05^95^. Significance testing for copy number variation was performed with Student’s T-test (two-sided), also with BH testing, but with a cut-off FDR of 0.005. Functional annotations and gene names were imported from prior analyses as described previously^96^. To identify the promoter site for Crz1 binding upstream of *FKS1*, we used Yeastract+^97^ to ascertain the presence of GNGGCKCA^98^ at contig PEKT02000002.1 position 1009439-1009439 (893-900 bases upstream from *FKS1* start) as the putative binding site for Crz1 (GAGGCCGCA). Contig edges were inferred using the B8441 v3 assembly, which demonstrates seven contigs^99^. To test for polyploidy, variants were re-called using GATK with the diploid flag to count heterozygous positions per 10 kb. Allele frequency counts were calculated from mpileups derived as above.

### Quantification of total chitin and β-glucan exposure

The chitin content in the cell walls of *C. auris* isolates and their segregants was compared using previously described methods^100^. Cells were grown in YPD at 30 °C for 5 h with or without 8 μg/ml anidulafungin, fixed with 50 mM thimerosal, and stained with 10 μg/mL Calcofluor-White (CFW) in the dark for 60 min. Stained cells were washed twice with PBS, and their fluorescence quantified using an Attune NxT flow cytometer. The plots represent three biological replicate experiments, in each of which 10,000 events were acquired. As a negative control, cells were treated as above but without the addition of CFW. Median Fluorescence Intensities (MFI) were determined using FlowJo v.10 software. The exposure of β-glucan at the *C. auris* cell surface was quantified by flow cytometry as described previously^101^. Cells were fixed overnight with thimerosal and stained with Fc-Dectin-1 and anti-human IgG linked to Alexafluor 488 (Jackson ImmunoResearch, Ely, UK). The fluorescence of 10,000 cells per condition was assayed using an Attune NxT flow cytometer. Median Fluorescence Intensity (MFI) was quantified using FlowJo v.10 software, and fold changes in β-1,3-glucan exposure were calculated relative to the control.

### Transmission electron microscopy

Transmission electron microscopy (TEM) was performed as described previously^101^. Cells were subjected to high pressure freezing and freeze substitution^102^, and fixed and stained with 1% osmium tetroxide and 0.5% glutaraldehyde. Ultrathin sections (60 nm) were prepared using lead citrate for contrast and imaged (JEOL 1400 JEM transmission electron microscope with ES1000W Gatan CCD camera). Cell wall sections were imaged at a nominal magnification of x100k. For each condition, approx. 30 cells were imaged and 10 measurements of inner and outer cell wall diameter taken for each cell using the line tool in ImageJ^103^.

### Cell wall carbohydrate analysis

Determination of cell wall mannan, chitin, and b-glucan content was achieved by acid hydrolysing the polymers, and quantifying mannose, glucosamine, and glucose content, respectively, by high-performance anion-exchange ion chromatography with pulsed amperometric detection (HPIC) as previously described^104^.

### Ploidy testing

The ploidy of cells in *C. auris* colonies growing on YPD plates containing 8 μg/ml anidulafungin was assayed by flow cytometry using previously described procedures^69^. Using a toothpick, cells were harvested directly from individual colonies and fixed overnight in 70% ethanol. The next day, cells were harvested by centrifugation (8,000 rpm, 5 min.) and resuspended in 50 mM sodium citrate, pH 7.5 and incubated with RNase A (250 µg per 10^7^ cells) and proteinase K (1000 µg per 10^7^ cells) for 4 h at 37 °C. Cells were then washed in PBS, resuspended in 0.25% Triton-X 100 (Sigma-Aldrich) and stained with SYBR Green I (1:500; Sigma-Aldrich) overnight at 4 °C overnight. Before flow cytometry, samples were sonicated and washed with PBS. Flow cytometry was performed on an Attune NxT flow cytometer using an excitation wavelength of 488 nm. SYBR Green I fluorescence was detected with a 530/30 band pass filter and 50,000 events, gated for single cells, were recorded for each sample. The proportions of individual cells that displayed ploidies of 1n, 2n, 3n, 4n or >5n were quantified using FlowJo v.10 software, using isogenic haploid and diploid colonies of *S. cerevisiae* grown on YPD plates as controls: W303-1B a (*MAT**a**, ade2, his3, leu2, trp1, ura3*); W303-1B 2M (*MAT**a**/MAT**α**, ade2/ade2, his3/his3, leu2/leu2, trp1/trp1, ura3/ura3*). The cells were also examined by fluorescence microscopy, staining them as described above with the addition of CFW (25 µg/ml) for 5 min. Cells were imaged using a DeltaVision Elite fluorescence microscope with a 60 x objective. Fluorescence excitation was generated by a LumencorLED light source and 10 μm Z-stacks of 50 images were captured by a pco.edge sCMOScamera. The 3D stacks were then deconvolved to remove out of focus light and maximum intensity projections used to create a 2D image by Image-J v1.54.

## Data availability

Raw reads have been deposited *via* SRI *via* BioProject accession PRJNA1373000.

## Acknowledgements

We acknowledge funding from the MRC Centre for Medical Mycology at the University of Exeter (MR/N006364/2, MR/V033417/1), MRC Doctoral Training Grants (MR/P501955/2, MR/W502649/1), Wellcome Trust Career Development Award (215239/Z/19/Z), Wellcome Trust Fellowship (219551/Z/19/Z), and the NIHR Exeter Biomedical Research Centre. The views expressed are those of the authors and not necessarily those of the NIHR or the Department of Health and Social Care. We thank the Exeter Sequencing Service facility and support from Wellcome Trust Institutional Strategic Support Fund (WT097835MF) to TB, NG, AB, and RAF, Wellcome Trust Multi User Equipment Awards (WT101650MA and 218247/Z/19/Z), Medical Research Council Clinical Infrastructure Funding (MR/M008924/1) and BBSRC LOLA award (BB/K003240/1). We thank the University of Exeter High-Performance Computing (HPC) facility, funded by the UK MRC Clinical Research Infrastructure Initiative (award number MR/M008924/1). BC and SM were funded *via* the St George’s Hospital Charity grant to TB (19-20-001). Medical Research Foundation Emerging Leaders award in AMR to TB (MRF-160-0009-ELP-BICA-0802), and the NIHR Exeter BRC. NG and AB also acknowledge the support of Wellcome Trust Investigator, Collaborative, Equipment, Strategic and Biomedical Resource awards (101873, 200208, 215599, 224323), and the MRC (MR/M026663/2, MR/Y002164/1). TB acknowledges salary support from MR/Y002164/1. We are grateful for comments on the manuscript from Dr Johanna Rhodes.

## Author contributions

All authors contributed to conceptualisation, writing and editing of the paper. TB originally conceived the study alongside AB, NG, and HG. TB and ST provided the clinical isolates and metadata for StG and K isolate series. All bioinformatic work was performed by HG with support and supervision from RF. BC and SM obtained comprehensive population profiling, colony morphotype analysis and EUCAST MIC testing, AP performed cell wall analysis, IL performed DNA extractions and microscopy, ME performed HPIC, TC performed additional experiments with DW.

## Notes

### Competing Interest Statement

TB has received speaker fees from Napp/ Mundipharma and Gilead Sciences, advisory board fees from Napp/ Mundipharma and research funding from Pfizer and Gilead sciences. All other authors declare no competing interests.

