## Supplementary information appendix for "*FKS1/2*-variant independent mechanisms underlying the emergence of resistance in echinocandin-refractory *Candida auris* infections"

### Supplementary note 1

In clinical isolates (StG1-5, K1-2), we also detected 178 synonymous variants, 42 insertions variants, 24 deletion variants, 24 intron-based variants, and 1 readthrough variant. We also identified three intergenic deletions (1-2 nt) across all clinical isolates located 664-1312 bases upstream of *FKS1*. Importantly, none of these mutations directly affected the putative Crz1 promoter binding site at 893-900 bases upstream but included GAA->G (664 bases upstream), CA->C (982 bases upstream) and TA->T (1,329 bases upstream). A further variable insertion (T->TAA-TAAAAAAAA, 1,527 bases upstream) was present in all clinical isolates except isolate StG2. These intergenic mutations may impact DNA conformation and interaction with promoter/binding sites, although this has specific interaction not been explored in *C. auris*. Only a single mitochondrial variant was present in the StG series, a 2-nucleotide insertion in isolates StG1-2 and StG3-4, with 11 separate small (1-3 base) intergenic substitutions/indels present across a series (isolates StG1-5: $n$ = 2, isolates K1-2: $n$ = 2) or across both ($n$ = 7).

Eight different nonsense mutations occurred exclusively in LCVs across the length of the gene, between amino acid position 68 and 494. Three different frameshift deletions between 5 and 23 bases also occurred exclusively in LCVs across the length of the gene, between amino acid positions 242-292, 367-463 and 516-531. Other mutations included R463S, C495F, and C507Y in LCVs and A516V and R531P in SCVs. *VPS5* also demonstrated significant variation in LCVs *vs* SCVs (58.8% *vs* 16.7%, adjusted *p*-value = $1.52 \times 10^{-2}$), with of up to three frameshift insertions of 11-97 bases in 9/17 LCVs, 1/7 clinical isolates and 2/18 SCVs.

We performed gene enrichment comparing those genes with significant variation between groups as defined above to the remaining genes using PFAM, KEGG and GO term annotation, GPI-anchor predictions, secreted protein and transmembrane domain predictions. After significance testing using Fisher's exact test using a Benjamini-Hochberg multiple testing cut-off of 0.05, We identified 76 PFAM, KEGG and GO terms driven by 24 loci that were enriched across the comparison between LCVs and SCVs (**Figure S5**). We did not identify any significantly enriched PFAM, KEGG and GO terms in the comparison between clinical and laboratory isolates. In addition to genes already discussed, this process highlighted variation in genes or associated intergenic regions for processes involving cell wall and membrane (*APS2*, *GDA1*, *TAZ1*), transcription and RNA processing (*IFH1*, *RAI1*, *STS1*, *UPT7*), DNA replication stress (*CAS1*, *LIA1*) and cellular morphology and cytoskeletal modelling (*OFD1*, *LOS1*, *PES1*, *CRN1*), adding to the candidate list for potential drivers of phenotypic variation in the comparison of LCVs and SCVs (**Table S4**).

Next, we compared copy number variation (CNV), using mean depth of coverage per gene locus across each genome in parent *vs* LCVs *vs* SCVs (**Figure S6A**). Intriguingly, the standard deviation for each locus was greater for LCVs and SCVs compared to clinical isolates (**Figure S6B**). There were ten loci with higher CNV in LCVs *vs* clinical isolates, and thirteen for SCV *vs* clinical isolates (**Figure S6C**). None of these areas of CNV reached above 1.98 normalised to genome-wide DOC, and all significantly different DOC were higher in LCVs/SCVs compared to clinical isolates. Six loci had higher CNV in both, including cell wall components *CSA1* and *PGA6*, plus transcription factors *BCR1* and *HP_3966* (**Table S5**). Four loci had higher CNV in LCVs, including cell-wall related *IFF4*, and regulators of morphology *NUS1* and *WOR2*. Seven loci had higher CNV in SCVs, including mitochondrial *COR1*, histone acetyltransferase *NTO1* and metabolic genes *ACS1*, *RIB7*, and *SPE4*. These variable loci provide candidates where CNV may play a role in tolerance and resistance, though this CNV does not appear to entirely explain the LCV/SCV phenotypic variation.

With regards to the elevated copy number across three orthologues of the Zorro3 retrotransposon encoding *FGR14* in clinical isolates, and an area of increased copy number in chromosome 1 featuring histone modifier *NTO1*: the *FGR14*-related CNV has been described[1] and even associated with multidrug resistance[2], but the presence of repetitive elements could yet reflect assembly errors. Other areas of CNV implied the loss of genes involved in transcription (*ARX1*, *IWR1*, *MRP49*), DNA maintenance (*CDC13*, *MRF1*, *PRD1*, *TMA17*), and histone deacetylation (*RCO1*), raising the question as to whether these areas of CNV represent gene losses that could contribute to the echinocandin tolerance/resistance.

We also examined the influence of sequencing of isolates grown directly from overnight cultures with anidulafungin by testing differences between mean normalised depth of coverage (DOC) for each locus by each group (student's T-test with Benjamini-Hochberg correction and false discovery rate (FDR) <0.005). There were no loci with significant differences in CNV between isolates sequenced directly on or off anidulafungin, consistent with the assumption that the genotype/phenotype of variants was preserved through generations.

The clear predominance of different *CRZ1* mutations in LCVs indicates a role for the calcium-calcineurin stress signaling pathway in resistance. Intergenic-associated mutations for *RCN2* were also present in LCVs. This pathway is implicated in echinocandin resistance mechanism development, given that the Crz1 transcription factor plays a role in cell wall remodeling e.g. *via* modulating *FKS* and *CHS* gene expression in *S. cerevisiae*[3,4].

Our findings of highly variable copy number across loci in LCVs/SCVs could be linked to the polyploid state, potentially including significant genome instability across isolates. The loci involved led us to hypothesise for roles in transcription factor and biofilm regulator *BCR1*, and cell wall genes components *CSA1* and *PGA6*, in resistance/tolerance. Furthermore, the morphology regulators *WOR2* and *NUS1*, which are implicated in *C. albicans* white-opaque switching and filamentation respectively[5,6], had higher CNV in LCVs. The higher CNV of metabolic, mitochondrial and histone acetyltransferase-related genes in SCVs also lends weight to the possibility that polyploidy-driven genome instability may confer drug resistance/tolerance through multiple mechanisms.

We used a range of techniques to explore the dynamic cell wall changes, including TEM and HPIC. We found that LCVs/SCVs had cell walls with an increased inner (glucan-chitin rich) cell wall diameter, while cells from LCVs exhibited a high glucan content, implying escape from glucan synthase inhibition in the face of anidulafungin drug pressure. Cell wall remodeling and chitin synthesis in *C. albicans* is regulated in part by *CRZ1* signaling[7]. Echinocandin treatment stimulated increases in cell wall chitin content *via* cell wall remodeling in *C. albicans*, analogous to our findings[8], with an elevated exposure of their pro-inflammatory β-glucan pathogen-associated molecular patterns (PAMPs)[9]. Echinocandin tolerance has long been associated with increased chitin content as a response to enable cell wall strengthening[10]. A recent solid-state NMR analysis of *C. auris* after caspofungin exposure indicated increases in cell wall strength through rigidification of mannan and β-1,3-glucan sidechains, rather than through increased chitin content[11]. We find that increased glucan content of cells from LCVs suggests an alternative adaptation mediated by *CRZ1* polymorphisms.

We considered the hypothesis that SCVs were petite mutants. Petite colony morphotype mutants are defective in mitochondrial respiration and may not confer echinocandin resistance but may prevent synergy of caspofungin with calcineurin inhibitors e.g. in *C. glabrata*[12]. SCVs have been described in *C. glabrata* in echinocandin microevolution experiments. These were associated with echinocandin resistance and preserved virulence, but with no apparent respiratory function deficiency[13]. We did not identify any mitochondrial genome mutations in our echinocandin resistant or tolerant sub-population, or any significant differences in intergenic variation between colony morphotypes. The petite phenotype is a respiration deficient variant demonstrating a small colony morphotype, that can be demonstrated through mitochondrial variants or genomic defects in oxidative phosphorylation[14] with reduced growth in YPD liquid or YPD-glycerol of *via* violet agar testing[15]. In *C. glabrata*, the petite phenotype does not confer resistance and has lower *in vivo* fitness[12] but may confer persistence in response to echinocandin exposure[16].

Several unexplored techniques to interrogate *C. auris* resistance/tolerance here include the transcriptome (to assess physiological difference) and epigenome (to understand chromatin changes and histone modifications). Investigating these avenues is supported by our data, which provide evidence of variation in transcriptional and chromatin/histone-related machinery. Epigenetic regulation of resistance in *C. auris* through chromatin modelling has not yet been fully evaluated, and this is an obvious next step in investigation following our findings. Roles for histone modification in *C. albicans* stress tolerance[17] and azole drug resistance[18] make it highly plausible such mechanisms could underlie resistance in *C. auris*.

#### Supplementary figures

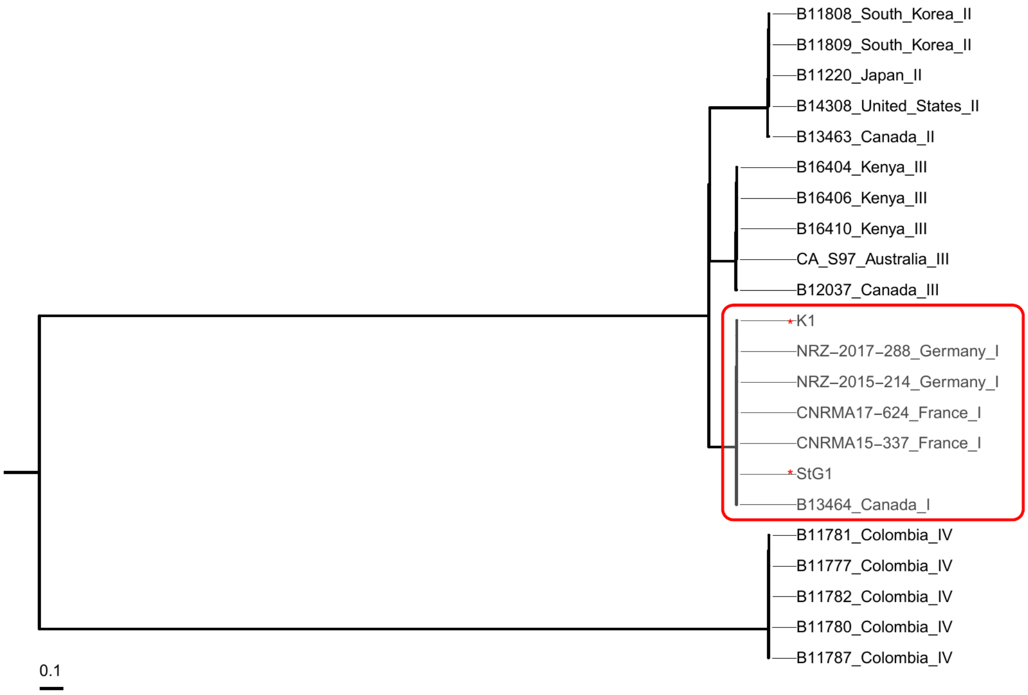


Figure S1: FastTree phylogeny demonstrating the placement of both parent clinical isolates (red asterisks) isolate StG1 (St George's Hospital) and isolate K1 (Kings College Hospital) and within clade I (red box). Isolates were randomly chosen as representatives of clades I-IV from a previous study[19]. Other isolates are designated by their isolate name, country of isolation, and known clade placement. Scale bar indicates number of substitutions per site.

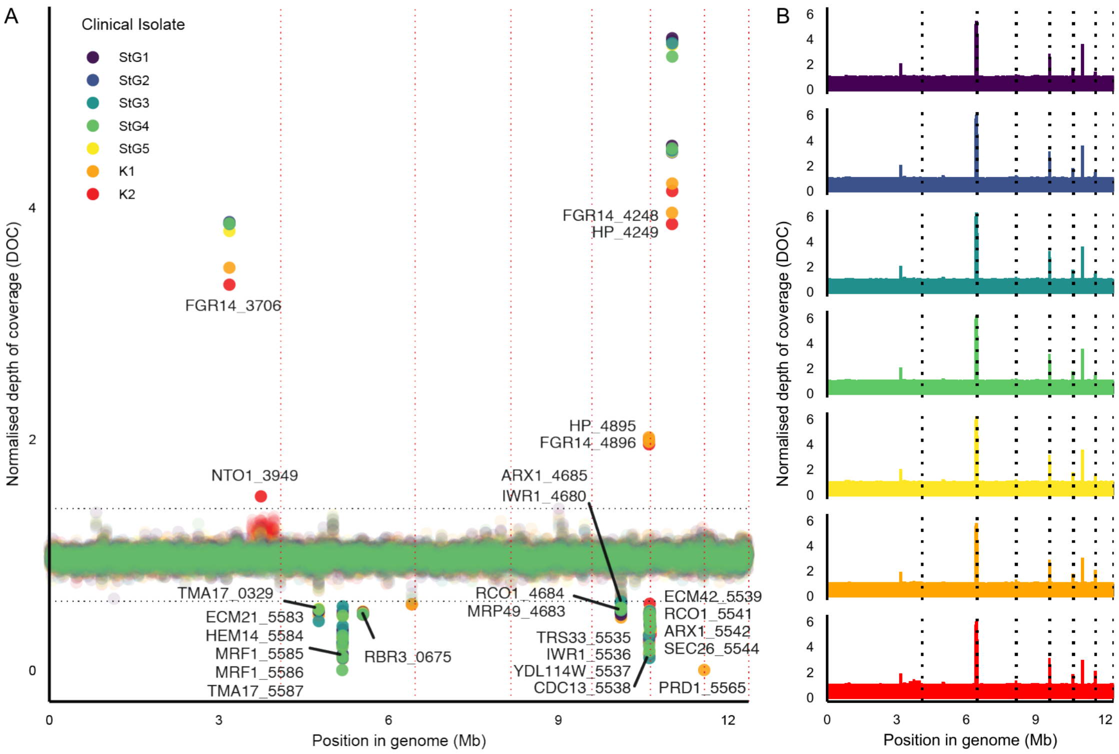


Figure S2: Clinical series copy number variation: **(A)** Copy number variation estimation by normalised depth of coverage per gene. Black horizontal line indicates cut-off (0.6/1.4), red vertical lines indicate contigs. **(B)** Normalised depth of coverage across the whole genome for each clinical isolate in 10 kb sliding windows. Red vertical lines indicate contigs. NSY: non-synonymous, SYN: synonymous.

152

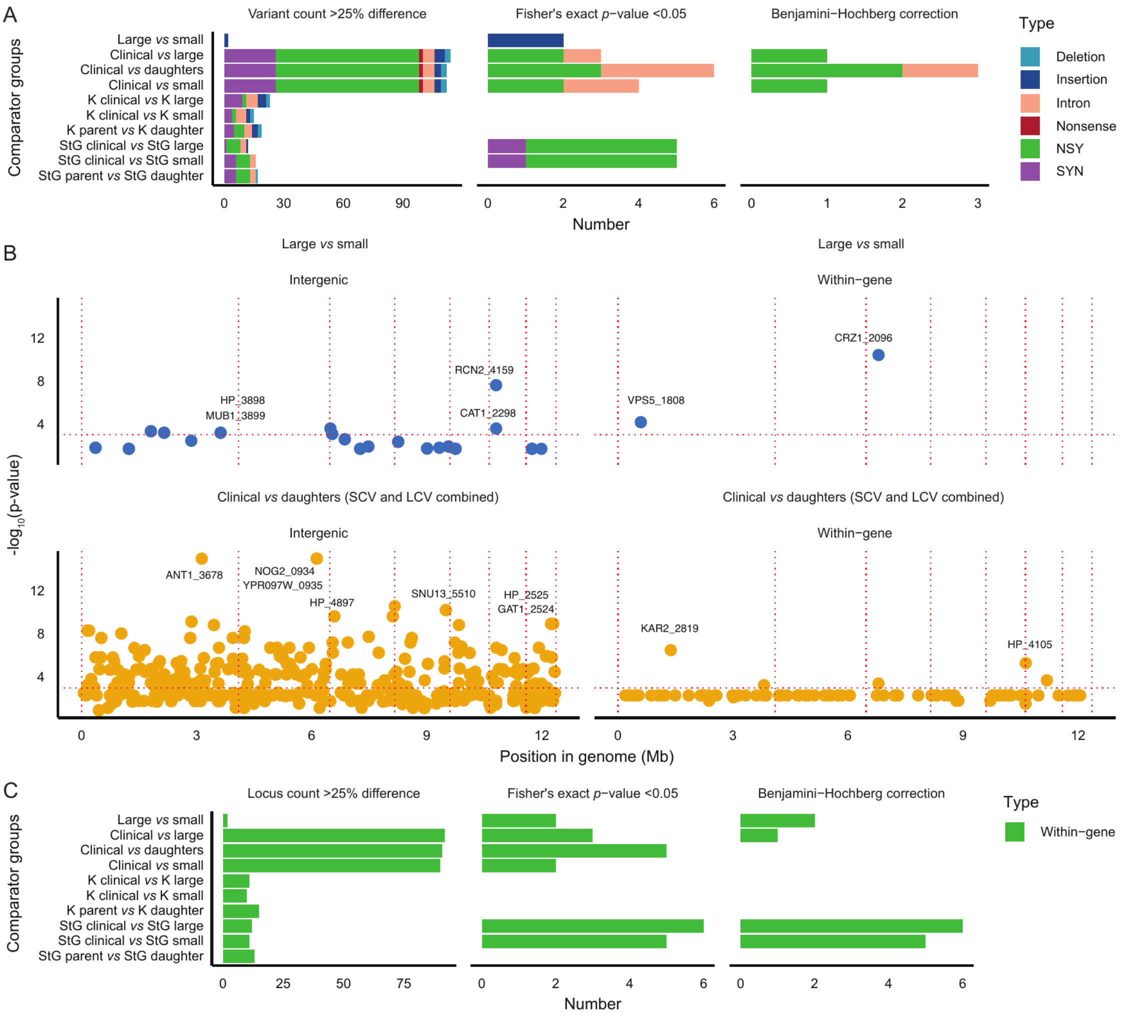


Figure S3: Genome-wide association: **(A)** Number of specific variants with >25% difference in proportion in between groups, tested for significance by Fisher's exact test with BH correction for within-gene variants. **(B)** Variable loci (genes with any variants) significant after BH correction plotted along genome position, with contigs marked in red. **(C)** Number of gene loci containing any variants with >25% difference in proportion, tested for significance as above. NSY: non-synonymous, SYN: synonymous.

153

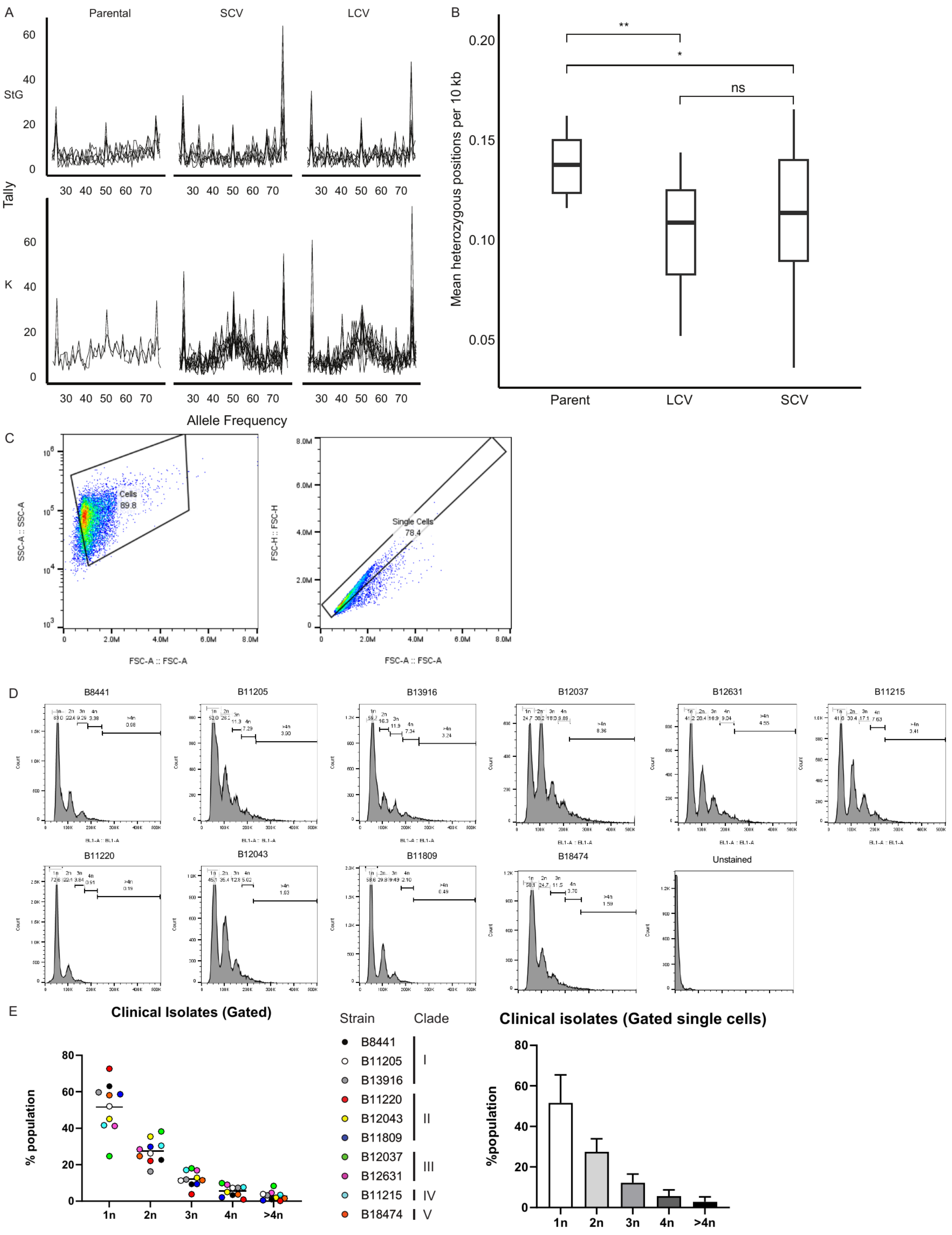


Figure S4: Bioinformatic and flow cytometric testing for polyploidy in *C. auris*. **(A)** Allele frequency tallies for each isolate, grouped by colony morphotype and clinical series. **(B)**

Numbers of heterozygous sites after variant calling with diploid flags. Boxplot indicates median, interquartile ranges as hinges, and whiskers extending to maximum value <1.5 times the interquartile range from the hinges. Significance levels indicate two-sided T-test, * = $p < 0.05$, ** = $p \leq 0.01$. **(C)** Single cell gating was performed by using the FSC-A vs FSC-H strategy. **(D-E)** Test isolates from epidemiologically derived clinical *C. auris* isolates, clades I-V[20], and frequencies of elevated ploidy.

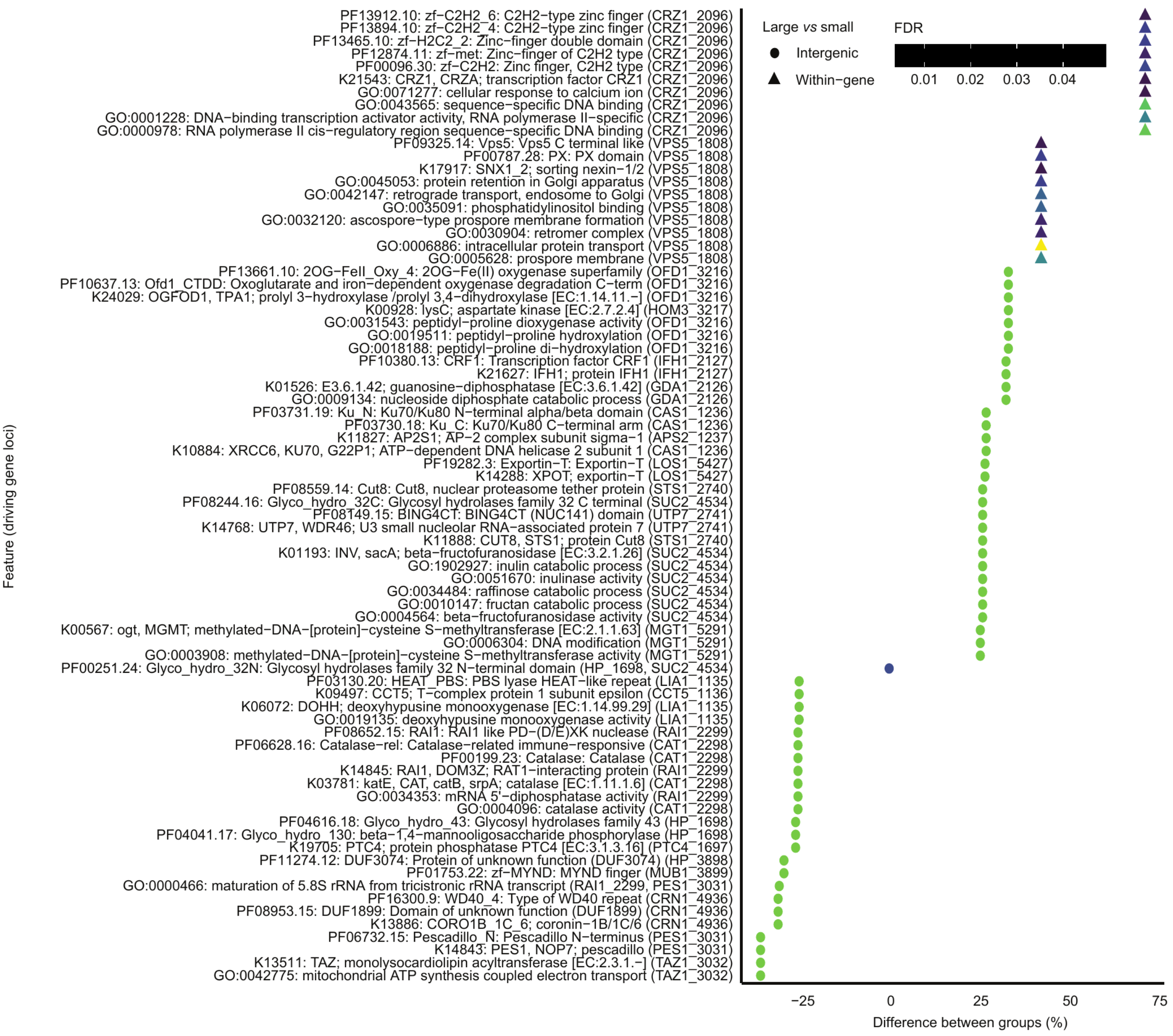

Figure S5: Loci annotated with GO terms, KEGG pathways and PFAM domains were tested for enrichment between LCVs and SCVs. Driving genes for each enriched term is indicated in brackets.

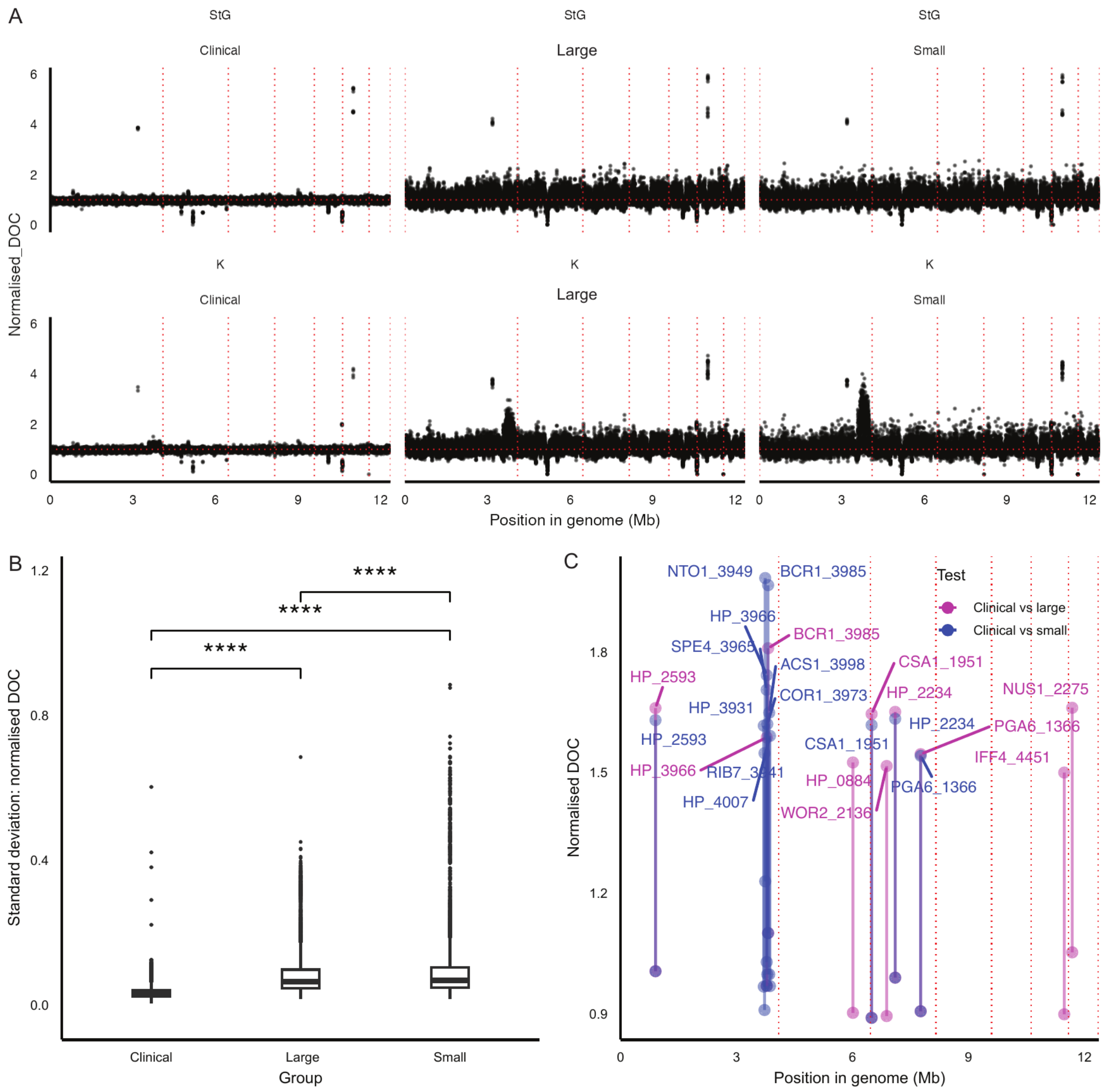


Figure S6: Copy number variation testing in morphotypic variant isolates: **(A)** Normalised depth of coverage (DOC) per gene locus for each isolate derived from StG and K clinical case series. **(B)** Standard deviation for each locus' normalised DOC (Wilcoxon test, significance level indicates p<0.0001). **(C)** Areas of CNV that were significantly different (>0.6 difference in normalised DOC, student's T-test with Benjamini-Hochberg correction and false discovery rate (FDR) <0.005) on comparison between large colony variants (LCVs) and small colony variants (SCVs) with clinical isolates.

#### Supplementary tables

| Gene | Stands for | Function in *C. auris* | Function in other human fungal pathogens |
|---|---|---|---|
| *BCK1_0735* | Bypass of C Kinase[21], member of Pkc1 cascade as mitogen-activated protein kinase kinase kinase (MAPKKK), cell wall stress stress response | Upregulated in response to caspofungin stress[22] | Deletion mutant in *C. albicans* is Caspofungin sensitive[23] |
| *BCY1_2818* | Bypass of cyclic AMP requirement[24] Protein kinase A regulatory subunit | Deletion associated with major loss of adhesion[25] and reduced virulence[26] | Deletion mutant in *C. albicans* has increased filamentation and white-opaque switching[27] |
| *CDC1_3971* | Cell division cycle protein[28] | - | Modification of the GPI anchor to facilitated glucan and cell wall protein binding across *Candida*[29] |
| *FGR14_3706* | Filamentous Growth Regulator[30] retroviral endonuclease-reverse transcriptase | Putative copy number variant[1] | Multiple putative functions involved in filamentation[30] |
| *GDI1_5447* | GDP-dissociation inhibitor[31] | - | Essential for secretory pathway and chitin deposition in *A. niger*[32] |
| *HAS1_4322* | Helicase Associated with Set1[33] involved in 18S rRNA biogenesis | - | Some evidence of variable expression in echinocandin resistant *C. albicans*[34] |
| *HP_3725* | Hypothetical protein | - | Orthologous to *C. albicans* translation initiation factor 33[35] |
| *HYR3_4100* | Hyphally regulated[36] GPI-anchored cell wall protein, also named Hil6[37] | Highly upregulated in murine catheter infection biofilm model *in vivo*[38]; upregulated in mature biofilms with concurrent echinocandin resistance[39] associated with efflux pump expression but also thought to be involved in extracellular matrix sequestration[40] and with signatures of selection in clades I, III and IV[41] | Mutations in areas of LOH in *C. albicans* serial clinical isolates with variable azole resistance[42] |
| *IWR1_4680* | Interacts With RNA polymerase II[43] putatively involved in nuclear protein import | - | - |
| *KAR2_2819* | KARyogamy 2 [@] Hsp70 family ATPase | - | Putative role in cell wall beta-glucan synthesis (GO:0070880) and ER translocation |
| *RBR3_0675* | Repressed by *RIM101*[44] cell wall gene | Mutations identified in clinical echinocandin resistance evolution[45], signatures of selection in clades I, III and IV[41] | Adhesin linked to glucan networks in *C. albicans*[46] |
| *SCF1_1458* | Surface colonisation factor1[25], also known as RBT1 | Surface colonisation factor1[25], required for adhesion and virulence and highly upregulated in murine catheter model *in vivo* biofilm[38] | Represented in network analysis of *C. albicans* gene expression network in caspofungin exposure[47] |
| *RRN3_3162* | Regulation of RNA polymerase I[48] | - | Understood to be upregulated in *C. albicans* on exposure to caspofungin[49], lost in fluconazole evolution experiment in *C. albicans*[50] |
| *SPT10_2077* | SuPpressor of Ty[51] Histone H3 acetylase | Upregulated in Amphotericin B resistant strains[52] | Putative target for antifungal therapy across *Candida* species[53] |
| *SVL3_1239* | Styryl dye Vacuolar Localization[54]; role in endocytosis | - | - |
| *VPS5_1808* | Vacuolar Protein Sorting 5[55] | - | - |

| | | | |
|---|---|---|---|
| *YPT6_3986* | Yeast Protein Two[55] endosome-to-golgi-mediating Rab family GTPase | - | Mutations can confer resistance to azole synergisers in several yeast species[56] |

Table S1: Variable loci in clinically evolved isolates, with gene name details, studies relating to function in *C. auris*, and role of orthologues in related *Candida* species. Gene names include final four number from B9J08_00 locus tags from the B8441 V2 reference strain.

| Gene | Stands for | Function in *C. auris* | Function in other human fungal pathogens |
|---|---|---|---|
| *ARX1_4685* | Associated with Ribosomal eXport complex[57] | Putative DNA-binding protein | Putative DNA-binding protein |
| *ARX1_5542* | ” | ” | ” |
| *CDC13_5538* | Cell Division Cycle[28] | - | Telomere binding and protecting across *Candida* sp.[58] |
| *ECM21_5583* | ExtraCellular Mutant[59] | - | Regulator of endocytosis, down-regulated in *C. albicans* in response to caspofungin[49], predicted to be involved in cell wall assembly with modelling of glucan synthesis gene expression networks and hypersensitivity to calcofluor white in mutant *S. cerevisiae*[60] |
| *ECM42_5539* | ” | - | Putative role in ornithine acetyltransferase/arginine biosynthesis, upregulated in amino acid starvation[61] |
| *FGR14_3706* | Filamentous Growth Regulator[30] retroviral endonuclease-reverse transcriptase | Putative copy number variant encoded by Zorro3 retrotransposase[1], part of a core orthologous group associated with azole resistance suggested to play a role in muti-drug resistance via regulation of transcription/RNA life-cycle as an RNA-dependent DNA polymerase[2] | Multiple putative functions involved in filamentation[30] |
| *FGR14_4248* | ” | ” | ” |
| *FGR14_4896* | ” | ” | ” |
| *HEM14_5584* | HEMe biosynthesis[62] | - | Upregulated in transcriptome of azole-exposed *C. glabrate*, thought to play a role in increased Erg11 activity[63] |
| *HP_4249* | Hypothetical protein | Putative zinc-binding transcription factor, identified as being associated with amphotericin B and fluconazole resistance[2] | - |
| *HP_4895* | ” | - | - |
| *IWR1_4680* | Interacts With RNA polymerase II[43] involved in protein import into nucleus | - | Nuclear import of RNA polymerase II in *S. cerevisiae*[64] |
| *IWR1_5536* | ” | - | ” |
| *MRF1_5585* | Mitochondrial peptide chain Release Factor[65] | - | Disruption leads to mitochondrial genome instability in *S. cerevisiae*[65] |
| *MRF1_5586* | ” | - | ” |
| *MRP49_4683* | Mitochondrial ribosome protein[66] | - | Component of large ribosomal subunit[66] |
| *NTO1_3949* | NuA Three Orf[67] | - | Putative histone acetyltransferase (HAT) complex subunit with role in chromatin remodeling and NuA3 binding[67] |

| | | | |
|---|---|---|---|
| *PRD1_5565* | PRoteinase yscD[68] | Putative proteinase more highly expressed in clade I isolate with lower *MDR1* expression[69] | Zinc metalloproteinase[70] thought to be upregulated in DNA replication stress[71] |
| *RCO1_4684* | Rpd3S histone deacetylase complex[72] | - | Increased acetylation of *FLO8* and *STE11* in mutant *RCO1* in *C. cerevisiae* |
| *RCO1_5541* | ” | - | ” |
| *SEC26_5544* | SECretory[73] | Downregulated in amphotericin B resistant strains[52] | Protein trafficking from ER to golgi in *S. cerevisiae*[73] |
| *TMA17_0329* | Translation Machinery Associated[74] | - | Deletion associated with chromosome instability[75] |
| *TMA17_5587* | ” | - | ” |
| *TRS33_5535* | Trapp (transport protein particle) Subunit[76] | - | Require for required for Ypt1-mediated autophagy in *S. cerevisiae*[77] |
| *YDL114W_5537* | - | - | - |

Table S2: Loci demonstrating copy number variation in clinical isolates.

| Strain name | Anidulafungin EUCAST MIC mode (range, µg/mL) |
|---|---|
| StG1 Parent | 2 (2-4) |
| StG1 SCV | 2 (2-4) |
| StG1 LCV | 8 (8-16) |
| K1 Parent | 1 (1-2) |
| K1 SCV | 1 (1-2) |
| K1 LCV | 4 (2-8) |

Table S3: EUCAST method antimicrobial testing for anidulafungin demonstrates elevated minimum inhibitory concentrations (MICs) in large colony variants (LCVs) compared to small colony variants (SCVs) and parent colonies derived from original clinical isolates.

| Gene | Stands for | Function in *C. auris* | Function in other human fungal pathogens |
|---|---|---|---|
| *ANT1_3678* | Adenine Nucleotide Transporter[78] | - | Small molecule transport at peroxisome membrane[78] |
| *APS2_1237* | clathrin Associated Protein complex Small subunit | - | Cell membrane protein sorting in *S. cerevisiae*[79] |
| *CAS1_1236* | Caspofungin sensitivity[80] | - | Disruption confers caspofungin hypersensitivity in *C. albicans*[80], also known as ORF 19.1135 and a Ku70 orthologue, shown to regulate telomerase/recombination and modulate telomere length and structure[81] |
| *CAT1_2298* | Catalase[82] | - | Deletion in *C. albicans* has higher sensitivity to neutrophil ROS production and lower virulence[82] |
| *CCT5_1136* | Chaperonin Containing TCP-1[83] | - | Role in actin cytoskeleton organisation[83] |

| | | | |
|---|---|---|---|
| *CRN1_4936* | Coronin[84] | - | Actin cytoskeleton component[84] |
| *CRZ1_2096* | Calcineurin-Responsive Zinc-finger protein[3] | Predicted to play a role in β-glucan masking, putatively increasing immune escape and virulence[85] | Transcription factor driving *FKS* expression in *S. cerevisiae*[3], coordinating a broad range of stress responses via calcium and calcineurin pathways, including cell wall remodelling[4] including in azole tolerance[86] and echinocandin tolerance in *C. albicans*[87,88]. Deletion in *C. glabrata* confers echinocandin susceptibility, thus therapeutic targeting has been suggested[89] |
| *GAT1_2524* | GATA family transcription factor[90] | - | Regulator of nitrogen source metabolism[90] |
| *GDA1_2126* | Guanosine DiphosphatAse[91] | - | Mutants defective in O-mannosylation in *C. albicans*[92] |
| *HOM3_3217* | HOMoserine requiring[93] | - | Amino acid synthesis[93] |
| *HOT13_4160* | Helper of TIM[94] | - | Modifier of Translocase of Inner Mitochondrial Membrane[94] |
| *HP_1698* | Hypothetical protein | - | - |
| *HP_2525* | Hypothetical protein | - | - |
| *HP_3898* | Hypothetical protein | - | - |
| *HP_4105* | Hypothetical protein | - | - |
| *HP_4897* | Hypothetical protein | - | - |
| *IFH1_2127* | Interacts with Fork Head[95] | - | Regulator of ribosomal protein transcription[96] |
| *LIA1_1135* | Ligand of eIF5A[97] | - | Understood to be expressed in DNA replication stress[71] |
| *LOS1_5427* | Loss Of Suppression[98] | - | Nucleoskeleton component in *S. cerevisiae*[99] |
| *MGT1_5291* | O-6-MethylGuanine-DNA methylTransferase[100] | - | DNA repair methyltransferase; protection from DNA alkylating damage[100] |
| *MUB1_3899* | MUlti Budding[101] | Deletion increases Rpn4 levels leading to higher azole efflux pump expression inc. Cdr1[102] | Critical factor in Rpn4 ubiquitylation and degradation[103] |
| *NOG2_0934* | NucleOlar G-protein[104] | - | GTPase involved in ribosome maturation[104] |
| *OFD1_3216* | Prolyl 4-hydroxylase-like 2-Oxoglutarate-Fe(II) Dioxygenase[105] | - | Modulates morphology by Ume6 stabilisation in *C. albicans*[106] |
| *PES1_3031* | Pescadillo homolog[107] | - | Mutants defective in virulence; thought to be involved in morphology and lateral yeast growth in tissues in *C. albicans*[107] |
| *PTC4_1697* | Phosphatase Two C[108] | - | Mutation confers sensitivity to azoles; thought to be involved in mitochondrial ion homeostasis[109] |
| *RAI1_2299* | Rat1p Interacting Protein[110] | - | RNA processing and degradation[110] |
| *RCN2_4159* | Regulator of CalciNeurin[111] | - | Also part of calcineurin stress pathway[111] |
| *SNU13_5510* | Small NUclear ribonucleoprotein associated [[112] | - | RNA processing in *S. cerevisiae*[112] |
| *STS1_2740* | Sec Twenty-three Suppressor[113] | - | Proteasome targeting[114] and ribosomal RNA stability/protein transport roles in *S. cerevisiae*[113] |
| *SUC2_4534* | SUCrose[115] | - | Invertase[116] |
| *TAZ1_3032* | TAfaZzin[117] | - | Cell membrane phospholipid cardiolipin remodelling[117] |
| *UTP7_2741* | U Three Protein[118] | - | Ribosomal biogenesis[118] |
| *YPR097W_0935* | - | - | Modifies ergosterol distribution, also known as Lipid-droplet Ergosterol Cortex (LEC1)[119] |

Table S4: Significantly variable loci and loci containing enriched terms between large (LCV) and small colony variants (SCVs). FDR: false discovery rate.

| Gene | Stands for | Function in *C. auris* | Function in other human fungal pathogens |
|---|---|---|---|
| *ACS1_3998* | Acetyl-CoA synthetase[120] | - | Implicated in *C. albicans* carbon source range and virulence[121] |
| *BCR1_3985* | Biofilm and Cell wall Regulator[122] | Glycerol-induced Filamentation-Competent factor, mutations causing variations in filamentation competence and enhances skin colonisation with improved fatty acid β-oxidation metabolism[123] | Transcription factor regulating cell wall synthesis and biofilm formation[122,124] |
| *COR1_3973* | CORe protein of QH2 cytochrome c reductase[125] | - | Mitochondrial inner membrane electron transport chain component[125] |
| *CSA1_1951* | *Candida* Surface Antigen[126] | Upregulated in *ex vivo* whole blood model of infection[127] | GPI-anchored, CFEM-domain, hyphally located, Bcr1 regulated cell wall protein[124] |
| *HP_0884* | Hypothetical protein | - | - |
| *HP_2234* | Hypothetical protein | - | - |
| *HP_2593* | Hypothetical protein | 24 nt deletion linked to pan-resistance strain[128] | - |
| *HP_3931* | Hypothetical protein | - | - |
| *HP_3966* | Hypothetical protein | - | Orthologous to transcription factor *Candida* Galactose gene Activator *CGA1*[129] |
| *HP_4007* | Hypothetical protein | - | - |
| *IFF4_4451* | Ipf family F[130] | Up-regulated in *C. auris* filamentation[131] and in ambisome-resistant strains[52], deletion has reduced adhesion[25], variants have been associated with echinocandin resistance in GWAS[132] | GPI-anchored, disruption reduces virulence/adherence in *C. albicans*[133] |
| *NTO1_3949* | NuA Three Orf[67] | - | Putative histone acetyltransferase (HAT) complex subunit with role in chromatin remodeling and NuA3 binding[67] |
| *NUS1_2275* | Nuclear Undecaprenyl pyrophosphate Synthase[134] | - | Regulation of filamentation, intracellular tracking and lipid homeostasis in *C. albicans*[6] |
| *PGA6_1366* | Predicted GPI-Anchored[135] | - | GPI-anchored[135] |
| *RIB7_3941* | Riboflavin biosynthesis[136] | - | Riboflavin synthesis across *Candida* species[137] |
| *SPE4_3965* | SPErmidine auxotroph[138] | - | - |
| *WOR2_2136* | White Opaque Regulator[139] | Under-expressed in *in vivo* catheter-associated biofilm[38] | Transcription factor implicated in *C. albicans* white-opaque switching[5] |

Table S5: Loci with higher copy-number variation in large (LCV) or small colony variants (SCV) compared to clinical isolates.
